# Brainstem *Dbh*^+^ Neurons Control Chronic Allergen-Induced Airway Hyperreactivity

**DOI:** 10.1101/2023.02.04.527145

**Authors:** Yujuan Su, Jinhao Xu, Ziai Zhu, Ji Chin, Le Xu, Haoze Yu, Victoria Nudell, Barsha Dash, Esteban A. Moya, Li Ye, Axel Nimmerjahn, Xin Sun

## Abstract

Chronic exposure of the lung to irritants such as allergen is a primary cause of asthma characterized by exaggerated airway constriction, also called hyperreactivity, which can be life-threatening. Aside from immune cells, vagal sensory neurons are important for airway hyperreactivity ^1–4^. However, the identity and signature of the downstream nodes of this adaptive circuit remains poorly understood. Here we show that a single population of *Dbh^+^*neurons in the nucleus of the solitary tract (nTS) of the brainstem, and downstream neurons in the nucleus ambiguous (NA), are both necessary and sufficient for chronic allergen-induced airway hyperreactivity. We found that repeated exposures of mice to inhaled allergen activates nTS neurons in a mast cell-, interleukin 4 (IL-4)-and vagal nerve-dependent manner. Single-nucleus RNA-seq of the nTS at baseline and following allergen challenges reveals that a *Dbh^+^* population is preferentially activated. Ablation or chemogenetic inactivation of *Dbh^+^* nTS neurons blunted, while chemogenetic activation promoted hyperreactivity. Viral tracing indicates that *Dbh^+^*nTS neurons, capable of producing norepinephrine, project to the NA, and NA neurons are necessary and sufficient to relay allergen signals to postganglionic neurons that then directly drive airway constriction. Focusing on transmitters, delivery of norepinephrine antagonists to the NA blunted allergen-induced hyperreactivity. Together, these findings provide molecular, anatomical and functional definitions of key nodes of a canonical allergen response circuit. The knowledge opens the possibility of targeted neural modulation as an approach to control refractory allergen-induced airway constriction.

---

Interoception, a concept first introduced over 100 years ago ^5^, defines a fundamental biological process whereby the nervous system senses and responds to the inner state of the body including that of internal organs ^6^. Recent work has begun to unravel the neural mechanisms for interoception in many tissues, including the larynx, heart, intestine, and bladder ^7–11^. However, knowledge remains limited on complete multi-synaptic circuits that sense signals from organs, integrate them in the nervous system, and respond to them through modulation of organ function.

The lung, given its large surface area, rich nerve innervation and direct exposure to aerosol environment, is a prominent source of interoceptive signals ^12–15^. In lung, airway constriction, triggered by exposure to irritants including allergen, is a normal protective physiologic response. It is driven by contraction of smooth muscles that wrap around the airways ^16^. Following chronic exposure, allergen would trigger exacerbated airway constriction, also termed airway hyperreactivity. The resulting restricted airflow is a primary asthma morbidity and a major cause of asthma-associated deaths.

It has been shown that in repeated allergen-challenged mice but not in naïve mice, vagotomy abolished airway hyperreactivity as assayed in the presence of methacholine, a synthetic acetylcholine analogue, suggesting dependence of the hyperreactivity on vagal reflex ^2^. Similarly, ablation or repression of *Trpv1^+^* or *Nav1.8^+^* vagal afferent neurons led to reduced airway hyperreactivity in allergen-challenged mice, but not methacholine-induced acute constriction in naïve mice ^3,4^. In the context of repeated allergen-induced airway hyperreactivity, the identity of the nodes downstream of vagal afferent neurons, their molecular signatures, and the nature of neurotransmitters and signals remain poorly defined.

A key knowledge gap is the brainstem neurons important for transmitting the chronic allergen signal. Mapping data from us and others show that lung-innervating vagal sensory neurons project exclusively to the nucleus of the solitary tract (nTS) region of the brainstem ^8,13,15,17^. Aside from lung, nTS integrates sensory inputs from many organs including the heart, gut and intestine ^8,9,14,18,19^. As a key signal relay center, neurons in the nTS project to higher order brain nuclei, as well as effector areas in the brainstem, including the dorsal motor nucleus of the vagus nerve (DMV) and nucleus ambiguous (NA) ^20–22^. Systematic analyses of the neuronal diversity of the nTS and their downstream nodes are critical for interrogating functional heterogeneity and specificity.

In this study, we dissect the mechanism of how normal lungs react to repeated allergen exposures, a disease-relevant adaptive response that is distinct from acute airway constriction. In wild-type mice, we show that repeated, but not single, allergen challenges led to localized activation of nTS neurons. This activation is reduced in either mast cell mutants, after treatment with interleukin 4 (IL-4) neutralizing antibody, or upon vagotomy. Genetic ablation of activated nTS neurons blunted allergen-induced airway hyperreactivity, but not goblet cell metaplasia or type 2 immune cell recruitment. Using single-nucleus RNA sequencing (snRNA-seq), we identified 13 distinct neuron types in the nTS. Using this dataset, we identified a single *Dbh^+^* sub-population that is enriched for allergen-activated neurons. Ablating or disrupting the function of *Dbh^+^*nTS neurons blunted airway hyperreactivity, while activating them in place of allergen exposure promoted hyperreactivity in sensitized airways. On the descending efferent side, *Dbh*^+^ nTS neurons project to parasympathetic neurons in the NA region of the brainstem. Chemogenetic inactivation of *Chat^+^* neurons in the NA, but not the DMV, attenuated airway hyperreactivity, while activation of NA neurons in place of allergen exposure promoted hyperreactivity in sensitized airways. Blocking norepinephrine receptors in the NA blunted allergen-induced airway hyperreactivity. Instead of projecting directly to lung, NA neurons project to postganglionic neurons in the trachea and extrapulmonary bronchi, which in turn target airway smooth muscles. Taken together, these *in vivo* data delineate a complete interoception neural circuit in the normal lung that is conditioned by repeated allergen exposures, and is composed of nodes in the brainstem that are both necessary and sufficient for allergen-induced airway hyperreactivity, mimicking exacerbated airway constriction in asthma.

## nTS neurons were activated upon repeated, but not single, inhaled allergen challenges

To investigate neurons involved in allergen response, we used a mouse model of asthma with chronic intranasal instillation of house dust mites (HDM), a known trigger for asthmatic responses in human and rodents ^23^. We employed a well-established regimen, with multiple doses of allergen administrations to lung (Fig. 1a) ^24,25^. This triggered the expected asthmatic responses including exacerbated airway smooth muscle constriction, also termed hyperreactivity, goblet cell metaplasia and type 2 immune cell infiltration (Fig. 1q, r, Extended Data Fig. 1a, 6d-f). Extending beyond previous findings indicating that allergen acts through vagal neurons ^3,4,26^, we addressed if allergen administered to lung could trigger brainstem neuron activation. Immunostaining for immediate early factor FOS, a neuronal activation marker, two hours after each of the challenges in serial sections of the whole brainstem revealed an increase of FOS^+^ cells in HDM-treated mice compared to saline-treated control mice (Fig. 1b, c, Extended Data Fig. 1b-v). Increased FOS^+^ cells were enriched in the dorsomedial part of the nTS region between Bregma -7.20 mm and Bregma -8.08 mm (Extended Data Fig. 1b-u) in comparison to other regions such as the adjoining area postrema (AP), DMV and hypoglossal nucleus (12N) (Extended Data Fig. 2a, b). This enrichment in the nTS is consistent with the findings that lung innervating sensory nerves project specifically to the same dorsomedial part of the nTS at these Bregma regions ^8,13,15,17^. The increase is found to be statistically significant only after the 4^th^ HDM challenge, but not after the previous three doses (Extended Data Fig. 3a-i), further corroborated by *Fos* transcript signal (Extended Data Fig. 4a, b). These findings suggest that there is conditioning of nTS neurons by repeated, but not single-dose administration of allergen.

**Fig. 1.**
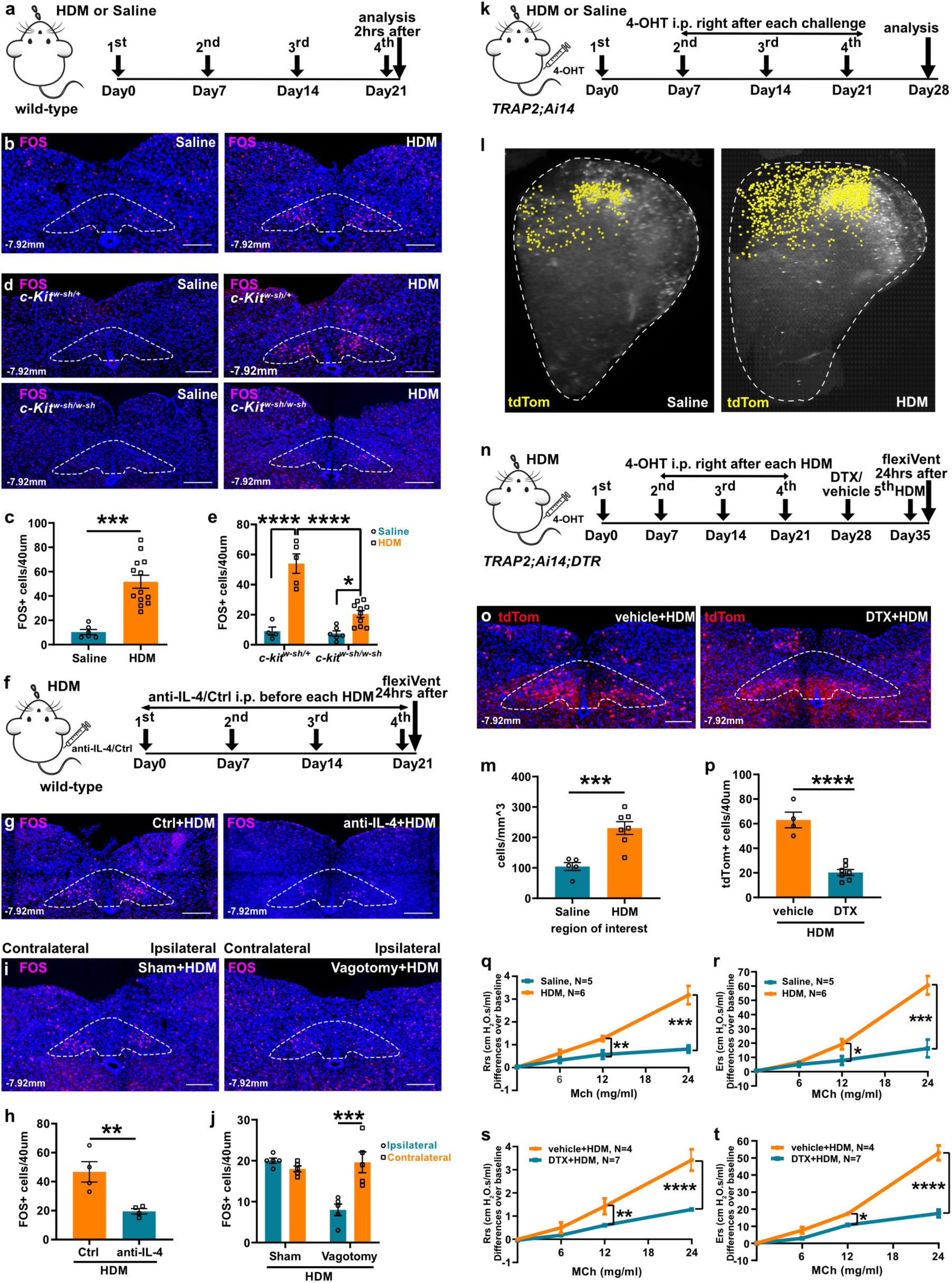
nTS neurons were activated upon repeated allergen challenges in lung. (**a**) Experiment scheme for HDM treatment in wild-type mice. (**b**, **c**) Representative FOS antibody staining (b) and quantification (c) showing increased FOS^+^ cells after 4^th^ HDM. Dashed circles outline nTS at the indicated Bregma level, as in subsequent related staining panels. Each data point represents the result from one animal, as in all subsequent graphs. (**d**, **e**) Representative FOS antibody staining (d) and quantification (e) showing decreased FOS^+^ cells in the nTS of *c-Kit^w-sh/w-sh^*, compared to *c-Kit^w-sh/+^* mice after HDM. (**f**) Experiment scheme for treatment with IL-4 neutralizing antibody in wild-type mice. (**g**, **h**) Representative FOS antibody staining (g) and quantification (h) showing decreased FOS^+^ cells in the nTS with anti-IL-4 neutralizing antibody treatment compared to isotype control antibody treatment. (**i**, **j**) Representative FOS antibody staining (i) and quantification (j) showing reduced signals on ipsilateral side with vagotomy, compared to contralateral side. (**k**) Experiment scheme for labeling allergen-activated neurons in *TRAP2; Ai14* mice. (**l**, **m**) Coronal view of CLARITY cleared hemisphere brainstem (l, dashed circles) and quantification (m) showing increased tdTomato^+^ cells after HDM. (**n**) Experiment scheme for ablating allergen-activated neurons in *TRAP2; Ai14; DTR* mice. (**o**, **p**) Representative nTS sections (o) and quantification (p) showing reduced tdTomato signals following DTX injection. (**q**, **r**) FlexiVent-measured maximal resistance (Rrs, q) and elastance (Ers, r) of the wild-type airway following increasing doses of MCh, demonstrating hyperreactivity after HDM challenge compared to saline control. (**s**, **t**) Maximal resistance (Rrs, s) and elastance (Ers, t) showing blunted airway hyperreactivity in DTX+HDM group of *TRAP2; Ai14; DTR* mice compared to vehicle controls. Statistical analysis was performed separately for each methacholine (MCh) concentration. Scale bars, 200 µm in b, d, g, i, o. Unpaired student’s t test was used for C, H, M and P, two-way ANOVA was used for e, j and q-t. Error bars represent means ± SEM. *p<0.05, **p<0.01, ***p<0.001, ****p<0.0001.

It is reported that mast cells are required for the development of allergen-induced airway constriction ^27–29^. Using the same mast cell deficient *c-Kit^w-sh/w-sh^* mice, we found a statistically significant decrease of FOS^+^ neurons in the nTS following HDM challenge compared to heterozygous *c-Kit^w-sh/+^* controls (Fig. 1d, e). These data suggest that signals relayed through mast cells are an important contributor to allergen-induced nTS activation.

Mast cells are among immune cells that produce type 2 immune signals, including interleukin 4 (IL-4), which are critical for allergen-induced airway hyperreactivity ^30,31^. It is reported that neutralization of IL-4 using antibodies can abrogate allergen-induced airway hyperreactivity ^30^. Using the same IL-4 neutralizing antibody and regimen, we found that administration of anti-IL-4, but not isotype control antibody significantly decreased the number of FOS^+^ neurons in the nTS (Fig. 1f-h).

To determine if vagal nerves are essential for allergen activation of nTS, we performed unilateral vagotomy before allergen administration. This led to a statistically significant decrease of FOS^+^ neurons on the ipsilateral operated side compared to the contralateral side following HDM challenge (Fig. 1i, j), suggesting that vagal nerves are required for transmitting allergen signals to the nTS.

To further quantify cumulative allergen-induced activation and also test a tool to manipulate these neurons, we crossed *Fos^2A-iCreER^*(*TRAP2*) mice to *Rosa-lxl-tdTomato* (*Ai14*) mice to label activated neurons with tdTomato. Following 4-OH tamoxifen injection after each of the HDM challenge or saline administration (2^nd^, 3^rd^ and 4^th^ doses, Fig. 1k), we cleared the whole brain and brainstem using CLARITY ^32^. We found a statistically significant increase of activated tdTomato^+^ neurons in the nTS of HDM-treated mice compared to saline-treated controls (Fig. 1l, m). This was further confirmed through serial sectioning and quantification (Extended Data Fig. 5a-c). These findings paved the way for manipulating these allergen-activated nTS neurons.

## Ablation of allergen-activated nTS neurons blunted airway hyperreactivity

To determine if activation of nTS neurons plays a role in allergen-induced responses, we crossed *Fos^2A-iCreER^; Rosa-lxl-tdTomato* (*TRAP2; Ai14*) mice to *Rosa-lxl-DTR* mice to express diphtheria toxin receptor (DTR) in allergen-activated neurons (*TRAP2; Ai14; DTR,* Fig. 1n). Following bilateral injection of diphtheria toxin (DTX) into nTS, we confirmed a decrease of allergen-activated neurons in the nTS (Fig. 1o, p).

To determine if targeted nTS neuron ablation affects airway hyperreactivity, we utilized a well-established assay that is tailored to test this exacerbated adaptive physiological response as a result of repeated allergen challenges ^1,33^. Following the last dose of HDM or saline administration, increasing doses of methacholine, a synthetic analogue of acetylcholine, was intratracheally administered into the airway to mimic what occurs in asthma patients, namely how acute triggers elicit chronically heightened airway exacerbation, the so-called “asthma attack”. Aside from directly acting on airway smooth muscles to trigger contraction, methacholine also signals to vagal afferent fibers to activate them which enhance smooth muscle constriction ^2–4,34^. In rodents, lung mechanics including airway hyperreactivity are commonly assayed using the flexiVent system.

Using this assay, in wild-type mice, as expected, we found that multiple doses of HDM sensitization and challenge led to further increase of flexiVent measurement of respiratory system resistance (Rrs) and elastance (Ers) compared to the saline control group, demonstrating allergen-induced airway hyperreactivity (Fig. 1q, r, N=5 for Saline, N=6 for HDM). As this hyperreactivity is abrogated by vagotomy ^2^, the differential airway constriction between the HDM and saline groups is primarily due to methacholine-activated vagal reflex rather than its direct effect on airway smooth muscles. Thus, by comparing the extent of airway constriction triggered by the same concentration of methacholine, we used the differences between experimental groups and corresponding controls as a measure of the impact of the chronically adaptive vagal circuit.

In the DTR-DTX experimental group, DTX ablation of allergen-activated nTS neurons led to blunted airway hyperreactivity compared to vehicle-injected group (Fig. 1s, t, N=4 for vehicle injection control, N=7 for DTX injection), suggesting that allergen-activated nTS neurons are part of the circuit conditioned by HDM to control airway hyperreactivity. In contrast, there was no change in HDM-induced goblet cell metaplasia, type 2 immune cell recruitment or expression of key type 2 cytokine genes *Il4*, *Il5* and *Il13* (Extended Data Fig. 6a-i). These data together suggest that while allergen-activated nTS neurons do not substantially modulate goblet metaplasia or type 2 immune recruitment, they are critical for amplifying allergen-induced airway hyperreactivity.

## Single nucleus transcriptomic signatures of the nTS allow dissection of diversity and selectivity

The nTS is known to receive diverse signals from many organs, including lung, heart, gut, intestine, etc ^8,9,14,18,19^. To identify which specific subset of the nTS neurons were activated by allergen response in lung, we started with defining the overall molecular diversity of the nTS through single-nucleus RNA sequencing (snRNA-seq) using flash frozen tissues. This method was chosen over single-cell RNA sequencing to avoid tissue digestion, cell dissociation and sorting which reduces neuronal viability and can also artificially induce cellular stress signatures ^35^. Using anatomical landmarks to minimize incorporation of the adjacent AP and DMV regions, we dissected the whole nTS region from adult wild-type mice (Fig. 2a). Transcriptome data from 12,966 nTS nuclei were obtained using 10x Genomics technology. Unsupervised clustering analysis revealed 8148 neurons and 4818 glial cells. Focusing on neurons, we further purified nTS neurons in silico by excluding AP and DMV neurons as identified by their profiles from two recently published snRNA-seq datasets of these regions ^36,37^. The remaining nTS neurons segregated into 13 clusters as shown in uniform manifold approximation and projection (UMAP) plots (Fig. 2b), a comparable number to a recent published nTS single cell-RNAseq dataset ^38^. All 13 neuronal clusters express pan-neuronal markers *Rbfox3* and *Snap25* (Fig. 2c, d), with little expression of glial marker genes (*GFAP*, *Olig1*, *Aqp4* or *Tmem119,* Extended Data Fig. 7a-d). Five neuronal clusters (Clusters 1, 2, 5, 9,12) are *Scl17a6^+^*glutamatergic excitatory neurons, while the remaining eight clusters (Clusters 3, 4, 6, 7, 8, 10, 11,13) are *Slc32a1^+^* GABAergic inhibitory neurons (Fig. 2e, f). Individual neuronal populations were clearly distinguished by a set of marker genes (Fig. 2g). We further validated the expressions of marker genes using UMAP plots and either our own RNAscope *in situ* hybridization assay (for excitatory clusters, Fig. 2h-q) or data from the Allen Brain Atlas (for inhibitory clusters, Extended Data Fig. 7e-l). A comprehensive analysis of genes encoding neurotransmitter/neuropeptide receptors, neurotransmitter synthesis/secretion machineries, neuropeptides and hormones revealed the molecular capabilities of each neuronal population in receiving and transmitting signals (Extended Data Fig. 7m-o).

**Fig. 2.**
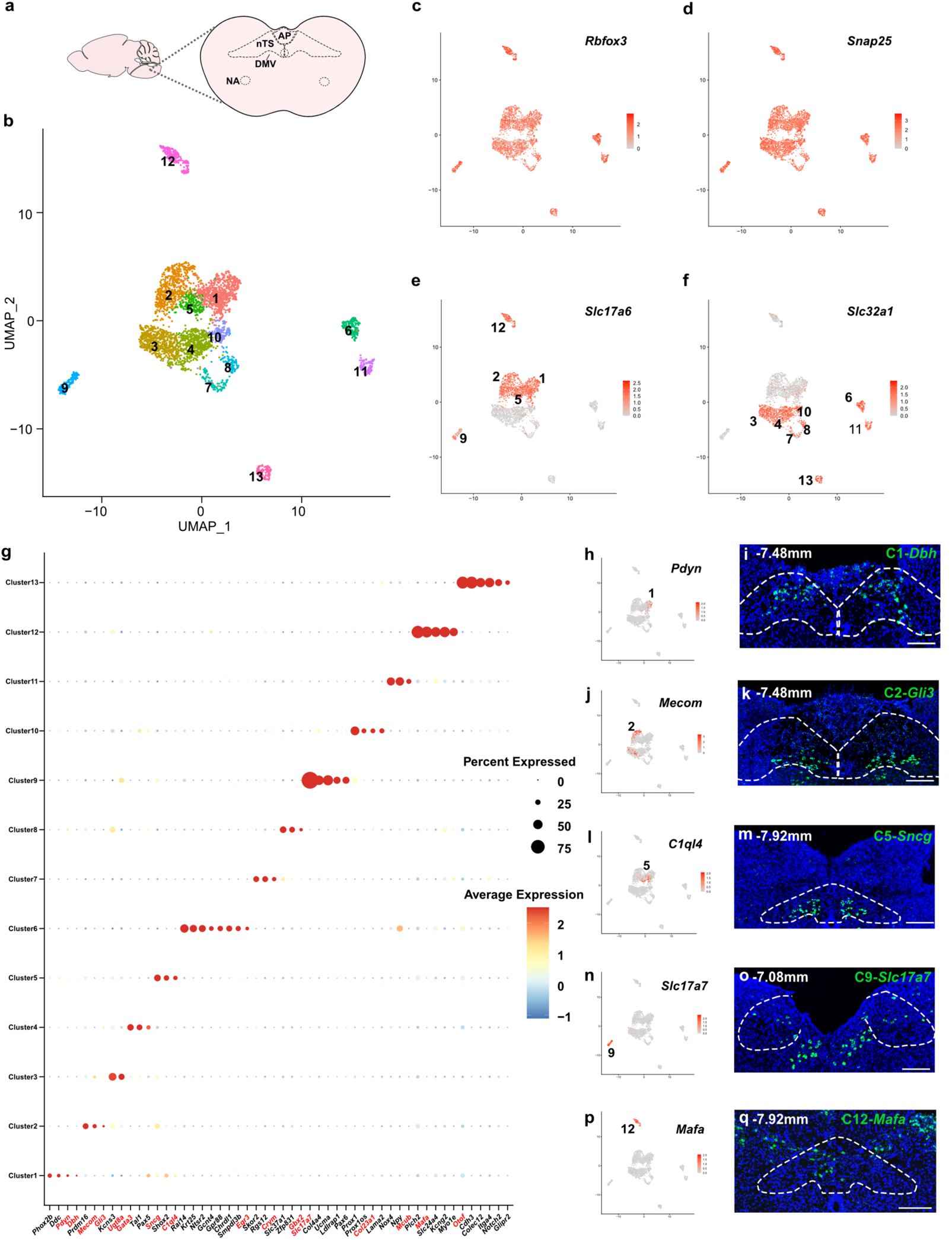
Single nucleus transcriptomic signatures of the nTS. (**a**) Diagram illustrating the relative locations of the nucleus of solitary tract (nTS), area postrema (AP), dorsal motor nucleus of the vagus (DMV) and nucleus ambiguous (NA). (**b**) Uniform manifold approximation and projection (UMAP) plots of snRNA-seq data showing 13 nTS neuron types. (**c-f**) UMAP plots showing expression of pan-neuronal markers *Rbfox3* (c) and *Snap25* (d) in all clusters, excitatory neuronal marker *Scl17a6* enriched in Clusters 1, 2, 5, 9 and 12 (e), inhibitory neuronal marker *Slc32a1* enriched in Clusters 3, 4, 6,7, 8, 10, 11 and 13 (f). (**g**) Dot plot showing top marker genes for each cluster. Gene names in red were used for validation by UMAP or RNAscope *in situ* hybridization as shown in Fig. 2h-q and Extended Data Fig. 7e-l. (**h-q**) UMAP plots (h, j, l, n, p) and RNAscope (i, k, m, o, q) showing expression of marker genes in the nTS excitatory clusters. Bregma levels shown are tailored to the layers with optimal signal for each marker. For i, k, m, o, q, outlines indicate nTS regions at the indicated Bregma level. Scale bars, 200 µm.

## *Dbh*^+^ neurons in the nTS were preferentially activated upon allergen challenge

To determine the molecular signature of allergen-activated nTS neurons, we compared nTS snRNA-seq signatures of wild-type mice 1.5 hours after the 4^th^ saline or 4^th^ HDM challenge. Compared to naïve nTS, the same 13 neuron subtypes were detected in saline- and HDM-challenged nTS (Fig. 3a, Extended Data Fig. 8a). Among the five clusters (Clusters 1, 6, 7, 10 and 12) that show Log2 average expression levels of *Fos* above 0, Cluster 1 is the only excitatory cluster that showed an increase of the percentage of *Fos*^+^ neurons (Fig. 3b, Extended Data Fig. 8b). Hence, we focused on Cluster 1 to determine its role in allergen-induced responses.

**Fig. 3.**
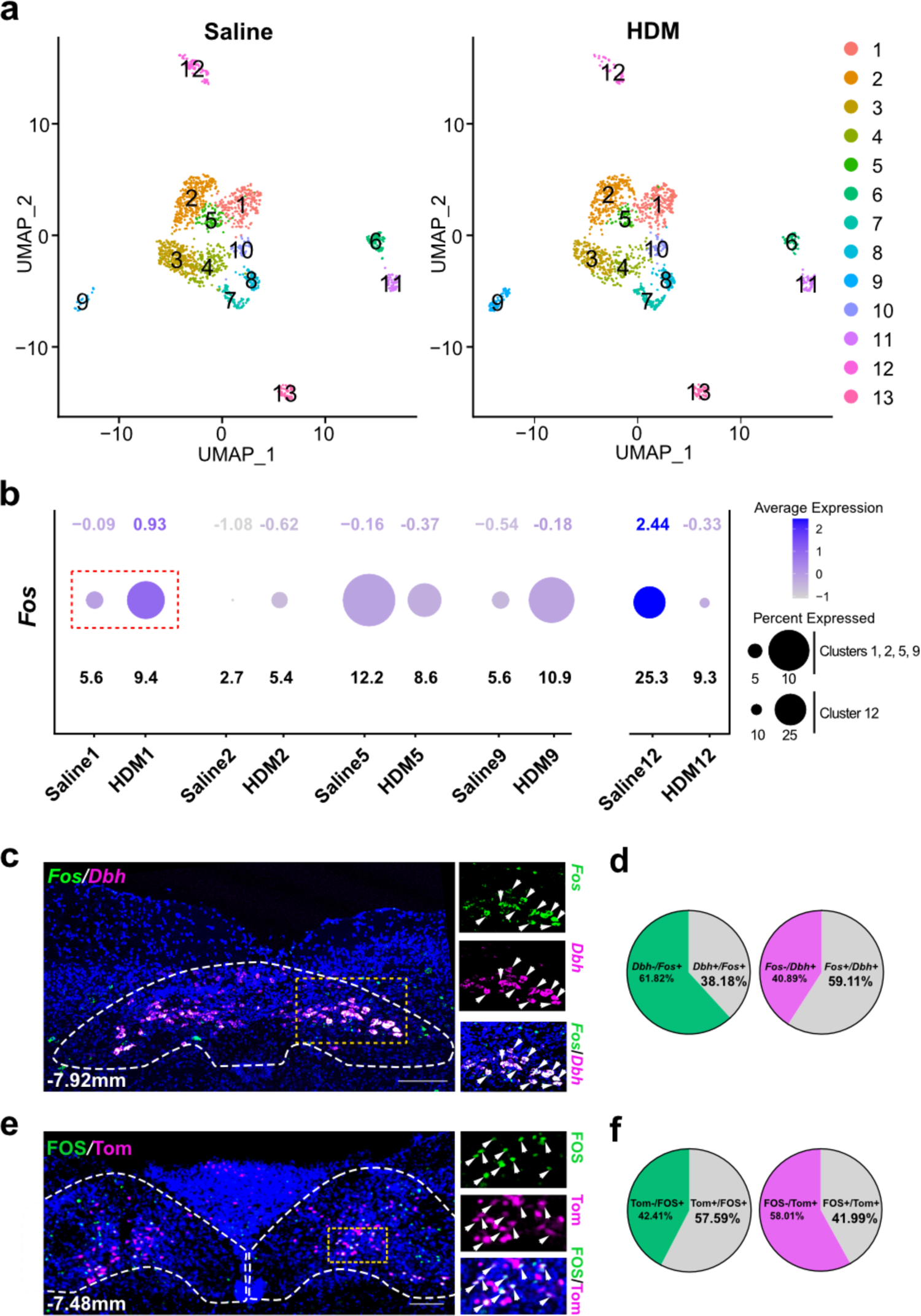
*Dbh^+^* neurons in the nTS were preferentially activated upon allergen challenge in lung. (**a**) UMAP plots showing the corresponding 13 neuron clusters in saline-(left) and HDM-challenged (right) nTS. (**b**) Dot plot of Fos expression in excitatory nTS clusters. Numbers on the upper row denote Log2 average expression levels, and numbers on the bottom row denote percentages of expressing cells. (**c**, **d**) Representative double *in situ* (c) and quantification (d) showing overlap between *Dbh* and *Fos* expression. (**e**, **f**) Representative FOS staining (e) and quantification (f) of *Dbh-cre; Rosa-lxl-tdTomato* (*Dbh-cre; Ai14*) nTS showing partial overlap between FOS and tdTomato. For c, e, outlines indicate nTS at the indicated Bregma, boxed areas magnified on the right, arrowheads indicate overlapped expression. Scale bars, 200 µm.

To validate that Cluster 1 neurons were activated upon HDM challenge, we performed double *in situ* hybridization with *Fos* and Cluster 1 marker gene *Dbh* (Fig. 3c). By *Dbh* RNAscope *in situ* hybridization on serial sections, we found that within nTS, *Dbh* expression is consistently detected between Bregma -6.96 mm and Bregma -8.08 mm, encompassing the regions where allergen-induced Fos^+^ cells were enriched (Extended Data Fig. 1b-u, Extended Data Fig. 9a). On individual sections, we observed a substantial overlap between *Dbh* and *Fos*, where 59.11% *Dbh^+^* neurons were *Fos^+^*, and 38.18% *Fos^+^* neurons were *Dbh^+^* (Fig. 3d, N=12 sections). To better assay the spectrum of allergen-activated neurons, we also screened a number of additional known nTS neuronal markers ^38–40^, including *Cnr1*, *Th* and *Cck*. We found that after HDM challenge, there were some *Fos* overlap with *Cnr1* and *Th*, but not *Cck* (Extended Data Fig. 10a-c). Among *Fos*^+^ neurons, the *Dbh^+^* neurons appear more enriched compared to *Cnr1*^+^ or *Th*^+^ neurons.

To further confirm that *Dbh*^+^ neurons were activated, and to validate a driver line to manipulate these neurons, we crossed transgenic *Dbh-cre* mice to *Rosa-lxl-tdTomato* (*Ai14*) mice. In *Dbh-cre; Ai14* mice, double *in situ* hybridization of *Dbh* and *tdTomato* showed >95% overlap of signals, confirming specificity of cre expression (Extended Data Fig. 11a, b, N=6 sections). After HDM administrations in *Dbh-cre; Ai14* mice, we observed a substantial overlap between tdTomato^+^ and FOS^+^ neurons in the nTS (41.99% tdTomato*^+^* neurons were Fos*^+^*, and 57.59% Fos*^+^* neurons were tdTomato*^+^*), further confirming that *Dbh*^+^ neurons were activated upon allergen challenges to lung (Fig. 3e, f, N=8 sections).

## Ablation or inactivation of *Dbh*^+^ neurons in the nTS blunted airway hyperreactivity

To determine if *Dbh^+^* nTS neurons are necessary for mediating allergen response, we employed three approaches. First, we performed chemical ablation by injecting anti-DBH antibody conjugated to the toxin Saporin (SAP). DBH-SAP was previously validated to specifically ablate DBH^+^ neurons with no effects on neighboring neurons ^41^. We injected anti-DBH-SAP stereotactically into both sides of the nTS of wild-type mice before allergen challenge (Fig. 4a). Compared to scrambled peptide-conjugated SAP (blank-SAP) control, anti-DBH-SAP group showed a clear reduction of DBH^+^ neurons after injection, confirming ablation efficiency (Fig. 4b). There is little change in the size of other excitatory nTS neuronal subsets as labeled by their specific markers, including *Gli3* (Cluster 2), *Sncg* (Cluster 5), *Slc17a7* (Cluster 9) or *Mafa* (Cluster 12), confirming ablation specificity (Extended Data Fig. 12a-f). Following HDM treatment, the anti-DBH-SAP injected group showed reduced FOS^+^ cells compared to blank-SAP control (Extended Data Fig. 13a-c). As a baseline control, in flexiVent assays, DBH^+^ neuron ablation had no effect on methacholine responses in naïve mice not exposed to HDM (Extended Data Fig. 13d-e, N=3 for blank-SAP+Saline, N=4 for anti-DBH-SAP+Saline). In contrast, following HDM challenge, the anti-DBH-SAP treated group showed statistically significant blunted airway hyperreactivity compared to the blank-SAP control group (Fig. 4c, d, N=4 for blank-SAP+HDM, N=9 for anti-DBH-SAP+HDM).

**Fig. 4.**
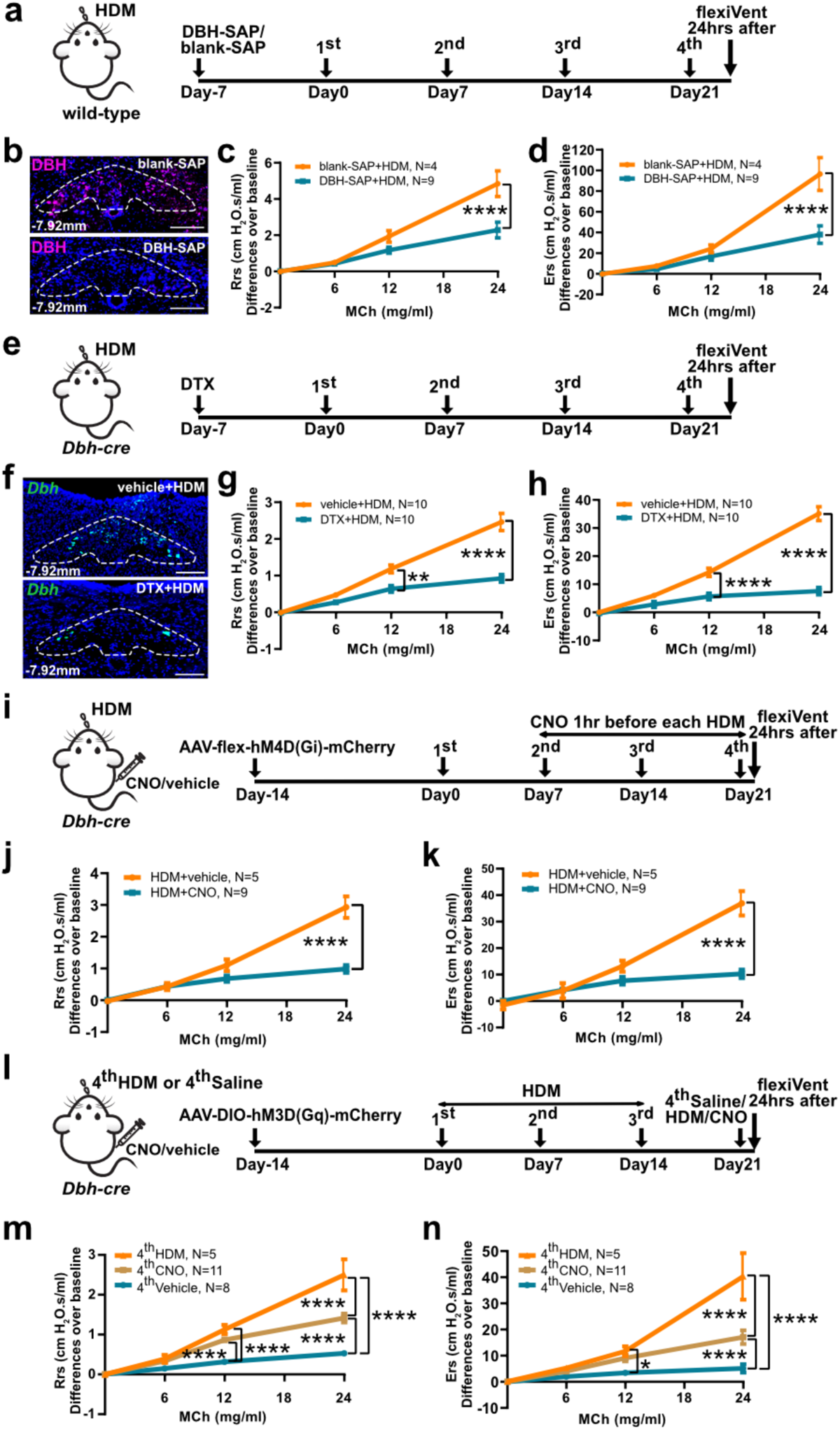
*Dbh^+^* neurons in the nTS mediates airway hyperreactivity. (**a**) Experiment scheme for chemical ablation of *Dbh^+^* nTS neurons. (**b**) Representative DBH antibody staining showing reduced DBH^+^ cells after ablation. Dashed circles outline dorsomedial part of nTS region at the indicated Bregma levels. (**c**, **d**) FlexiVent data showing blunted airway hyperreactivity after DBH-SAP. (**e**) Experiment scheme for genetic ablation of *Dbh^+^* nTS neurons. (**f**) Representative *Dbh in situ* hybridization showing reduced expression after DTX. (**g**, **h**) FlexiVent data showing blunted airway hyperreactivity after DTX. (**i**) Experiment scheme for chemogenetic inhibition of *Dbh^+^* neurons. (**j**, **k**) FlexiVent data showing blunted airway hyperreactivity after CNO injection. (**l**) Experiment scheme for chemogenetic activation of *Dbh^+^* neurons. (**m**, **n**) FlexiVent data showing increased airway hyperreactivity after CNO, in place of the 4^th^ HDM. All groups received the 1^st^-3^rd^ HDM challenge. Scale bars, 200 µm in b, f. Two-way ANOVA was used for c, d, g, h, j, k, m, n. Statistical analysis at each MCh concentration was performed separately. Error bars represent means ± SEM. *p<0.05, **p<0.01, ****p<0.0001.

Second, we performed genetic ablation by crossing *Dbh-cre* mice to *Rosa-lxl-DTR* mice (*Dbh-cre; DTR*) and then injected DTX bilaterally into nTS (Fig. 4e). This led to efficient loss of *Dbh*^+^ neurons in the nTS (Fig. 4f). Following this targeted genetic ablation, we carried out HDM challenge and found that allergen-induced airway hyperreactivity was blunted in the DTX group compared to vehicle-injected control group (Fig. 4g, h, N=10 for vehicle+HDM, N=10 for DTX+HDM).

Third, we performed chemogenetic inactivation using Designer Receptors Exclusively Activated by Designer Drugs (DREADD) system. We injected AAV-flex-hM4D(Gi)-mCherry bilaterally into nTS of *Dbh-cre* mice, eliciting efficient and specific expression in the nTS, but not nearby regions such as AP (Fig. 4i, Extended Data Fig. 16a, b). Following robust expression, we started the HDM regimen and administered Clozapine-N-Oxide (CNO) to activate hM4D(Gi) before the 2^nd^ to the 4^th^ HDM, sparing the first HDM to allow for sensitization of the immune system which is required for subsequent activation of the nervous system ^3^. We found that the CNO group showed blunted airway hyperreactivity response compared to vehicle group (Fig. 4j, k, N=5 for HDM+vehicle, N=9 for HDM+CNO). Taking together, data from chemical ablation, genetic ablation and chemogenetic inactivation demonstrate that *Dbh*^+^ nTS neurons constitute an essential node of the circuit critical for allergen-induced airway hyperreactivity.

## Activation of *Dbh*^+^ neurons in the nTS induced airway hyperreactivity

To address if *Dbh*^+^ neurons in the nTS are sufficient for mediating allergen response, we used DREADD targeted activation. We injected AAV-DIO-hM3D(Gq)-mCherry bilaterally into nTS of *Dbh-cre* mice, eliciting efficient and specific expression in the nTS (Fig. 4l, Extended Data Fig. 17a). Following this robust expression, we started the HDM regimen and administrated CNO in place of the 4^th^ HDM challenge while all groups received the 1^st^ to the 3^rd^ doses of HDM. The flexiVent data showed that while not to the full extent as HDM, CNO activation of *Dbh^+^* neurons in place of the 4^th^ dose was sufficient to induce increased airway hyperreactivity compared to saline control (Fig. 4m, n, N=5 for 4^th^ HDM, N=11 for 4^th^ CNO, N=8 for 4^th^ Vehicle). In contrast, in naïve mice not exposed to HDM, CNO activation (the 4^th^ dose) of *Dbh^+^*neurons had no effect (Extended Data Fig. 17b-d, N=3 for 4^th^ Saline+vehicle, N=5 for 4^th^ Saline+CNO). Taking both loss- and gain-of-function data together, these findings suggest that the activity of *Dbh*^+^ neurons in the nTS is both necessary and sufficient for mediating allergen-induced airway hyperreactivity.

In none of the three loss-of-function experiments, nor the chemogenetic gain-of-function experiment did we observe any changes in HDM-induced goblet cell metaplasia, increased immune cell infiltration or expression of *Il4*, *Il5* and *Il13* (Extended Data Fig. 14a-f, 15a-f, 16c-i, 18a-h). These findings together confirm findings from *TRAP2; Ai14; DTR* mice (Extended Data Fig. 6a-i). We also assayed for possible effects on other aspects of lung function. Following either HDM challenge or CNO-activation, compared to saline controls, we found no significant difference in minute ventilation, respiratory frequency, tidal volume, or metabolic rate, as measured by plethysmography (Extended Data Fig. 19a-g). This is consistent with previous report that HDM challenge in mice did not change respiratory parameters ^42^. These results indicate that CNO activation of *Dbh^+^* nTS neurons induced airway hyperreactivity in sensitized airways without affecting respiration.

## Parasympathetic neurons in the NA are downstream of *Dbh^+^* nTS and upstream of airway-innervating postganglionic neurons

To map the downstream targets of *Dbh*^+^ nTS neurons and their role in mediating allergen-induced airway hyperreactivity, we injected AAV-flex-tdTomato to the nTS of *Dbh-*cre mice (Fig. 5a). No fiber was detected in either the trachea or the lung, indicating that *Dbh*^+^ nTS neurons do not innervate the airway directly. We then sectioned and screened the whole brainstem and found that tdTomato^+^ fibers project to the NA which is composed of *Chat^+^* neurons (Fig. 5b). Our data suggest that *Dbh*^+^ nTS neurons project to the NA.

**Fig. 5.**
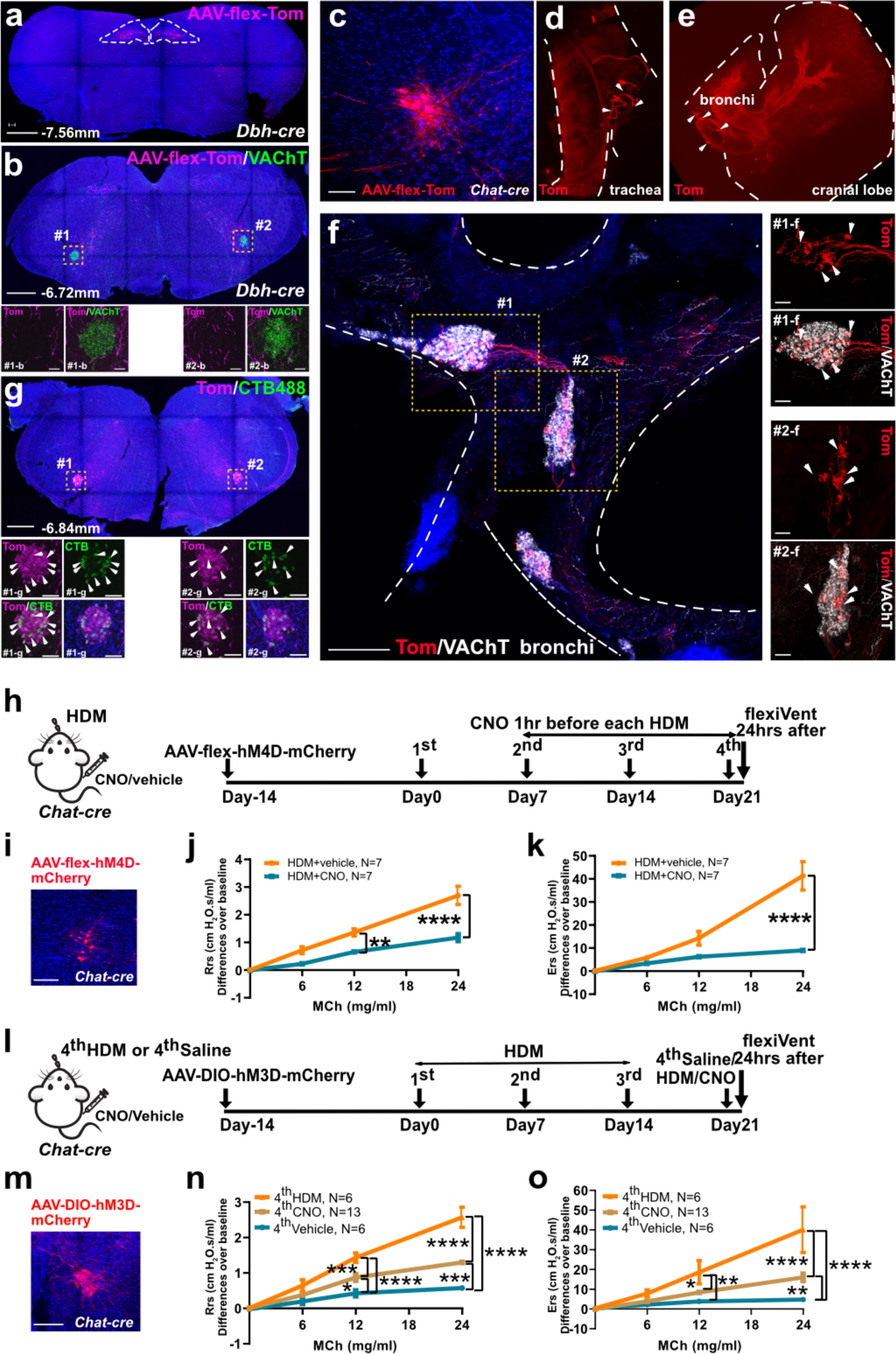
Parasympathetic neurons in the NA downstream of the nTS are necessary and sufficient for allergen-induced airway hyperreactivity. (**a**) Representative brainstem section at indicated Bregma level showing specificity of the injection of AAV-flex-tdTomato into bilateral nTS (outlined) of *Dbh-cre* mouse. (**b**) In the same mouse shown in A, tdTomato^+^ nerves project to NA region (VAChT^+^). Boxed areas are magnified below. (**c**) Injection of AAV-flex-tdTomato into NA of *Chat-cre* mouse. (**d**, **e**) In the same mouse shown in c, light sheet images showing tdTomato^+^ fibers innervate the trachea (d, arrowheads) and extrapulmonary bronchi (e, arrowheads). (**f**) Confocal stack (300 µm) of extrapulmonary bronchi showing NA-originated tdTomato^+^ fibers innervate postganglionic parasympathetic ganglia (VAChT^+^, arrowheads). Boxed areas are magnified on the right. (**g**) In *Chat-cre; Ai14* mice, injection of CTB488 in dorsal trachea led to overlapping CTB488^+^ and tdTomato^+^ signals in the NA (arrowheads). (**h**) Scheme for chemogenetic inhibition of NA *Chat^+^* neurons. (**i**) AAV-flex-hM4D-mCherry signals in the NA of *Chat-cre* mouse. (**j**, **k**) FlexiVent measurements indicate blunted airway hyperreactivity after CNO injection. (**l**) Scheme for chemogenetic activation of NA *Chat^+^* neurons. (**m**) AAV-DIO-hM3D-mCherry signals in the NA of *Chat-cre* mouse. (**n**, **o**) FlexiVent measurements indicate increased airway hyperreactivity after CNO, in place of the 4^th^ HDM. Scale bars, 100 µm in c, i, m, 200 µm in f (50 µm in magnified views), 500 µm in a, b (50 µm in magnified views) and g (100 µm in magnified views). Two-way ANOVA analysis for j, k, n and o were performed separately at each MCh concentration. Error bars represent means ± SEM. *p<0.05, **p<0.01, ***p<0.001, ****p<0.0001.

To address if NA neurons project directly to lung, we first stereotaxically injected AAV-flex-tdTomato to the NA of *Chat-cre* mice (Fig. 5c). Both light-sheet imaging of intact trachea/bronchi/lung (Fig. 5d, e) and confocal imaging of thick sections (Fig. 5f) revealed that NA-originated tdTomato^+^ fibers project to postganglionic parasympathetic ganglia residing in both the trachea and the bronchi, rather than directly to smooth muscle cells in lung (Extended Data Fig. 20a-e). In comparison, stereotaxic injection of AAV-flex-Tom into the DMV and the nearby 12N of *Chat-cre* mice labeled fibers that passed by the space between the trachea and the esophagus without innervating the trachea or the bronchi (Extended Data Fig. 20f, g).

To confirm NA to postganglionic neuron connection from the retrograde direction, we introduced retrograde tracing dye CTB488 dorsal to the trachea, where postganglionic parasympathetic ganglia are enriched. In whole brainstem sections, we found CTB488 labeling of *Chat^+^* neurons in the NA region, with little cell body labeling in other regions including the DMV and the 12N (Fig. 5g).

To directly address the role of *Chat*^+^ neurons in the NA in mediating allergen-induced airway hyperreactivity, we utilized the DREADD system. To inactivate *Chat^+^*neurons in the NA, we injected AAV-flex-hM4D(Gi)-mCherry bilaterally into NA of *Chat-cre* mice (Fig. 5h, i). We found that the CNO group showed blunted airway resistance and elastance responses compared to the vehicle-injected control group (Fig. 5j, k, N=7 for HDM+vehicle, N=7 for HDM+CNO).

Conversely, to specifically activate *Chat^+^* neurons in the NA, we injected AAV-DIO-hM3D(Gq)-mCherry bilaterally into NA of *Chat-cre* mice (Fig. 5l, m). While not as potent as HDM, activation of *Chat^+^* NA neurons by CNO is sufficient to induce increased airway hyperreactivity compared to saline (Fig. 5n, o, N=6 for 4^th^ HDM, N=13 for 4^th^ CNO, N=6 for 4^th^ Vehicle). In naïve mice not exposed to HDM, we found that CNO activation of *Chat^+^*NA neurons had no added effect on airway constriction under increasing doses of methacholine (Extended Data Fig. 21a-c, N=3 for 4^th^ Saline+vehicle, N=6 for 4^th^ Saline+CNO).

Previous studies have shown that DMV, a key node of autonomic control ^43^, is also a target of nTS neurons ^43,44^. To test if DMV may also act downstream of *Dbh^+^* nTS neurons in the allergen circuit, we performed DREADD-based inactivation or activation of *Chat^+^* neurons in the DMV (Extended Data Fig. 22a, d). In contrast to results from the NA, these DMV perturbations showed little effect in modulating HDM-induced airway hyperreactivity (Extended Data Fig. 22b, c, e, f).

Our data together indicate that *Chat^+^* neurons in the NA are necessary and sufficient in mediating allergen-induced airway hyperreactivity.

## Single nucleus transcriptomic signatures of the NA allow dissection of allergen signaling relay

To determine the molecular nature of the signaling relay leading to NA activation, we utilized snRNA-seq to define NA neuron signatures. As all NA neurons express *Chat*, in the brainstem of *Chat-cre; CAG-Sun1/sfGFP* mice, we collected around the small GFP^+^ region that corresponds to NA, and avoided DMV and 12N regions which are also GFP^+^. Among all the neurons identified in the snRNAseq dataset, we focused on the 190 *Chat^+^*neurons, which segregated into 3 clusters as shown in the UMAP plot (Fig. 6a). Individual neuronal populations were distinguished by marker genes (Fig. 6b), which were verified by UMAP plots and caudal brainstem *in situ* hybridization data from the Allen Brain Atlas (Fig. 6c-h). Our neuron number and neuron type clusters are comparable to the two recent published NA single cell datasets ^45,46^.

**Fig. 6.**
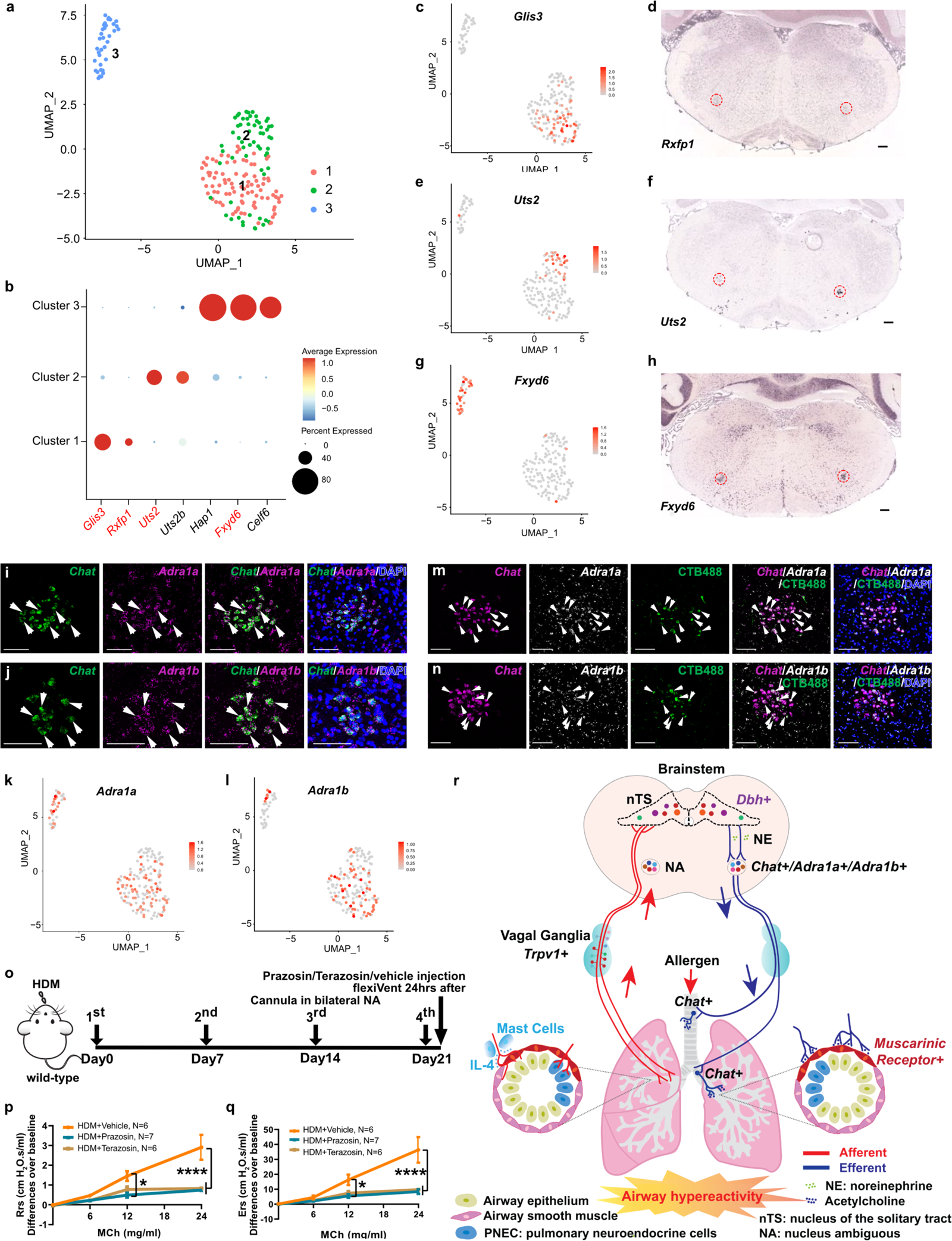
Blocking norepinephrine receptors in the NA blunted allergen-induced airway **hyperreactivity.** (**a**) UMAP plot of NA snRNA-seq data showing 3 NA neuron types. (**b**) Dot plot showing top marker genes for each cluster. Gene names in red were used for validation by UMAP plots or *in situ* hybridization as shown in c-h. (**c**-**h**) UMAP plots (left) and *in situ* hybridization images from Allen Brain Atlas (right) showing expression of marker genes. (**i**, **j**) Double *in situ* hybridization showing *Chat^+^* neurons in the NA express norepinephrine receptors *Adra1a* and *Adra1b* (arrowheads). (**k**, **l**) UMAP plots showing expression of *Adra1a* (k) and *Adra1b* (l) in all NA clusters. (**m**, **n**) Injection of CTB488 to dorsal trachea in wild-type mice labeled neurons in the NA that express *Chat* and *Adra1a* (m) and *Adra1b* (n) by RNAscope *in situ* hybridization (arrowheads). (**o**) Experiment scheme for norepinephrine receptor antagonist treatment. (**p**, **q**) FlexiVent data showing decreased hyperreactivity after Prazosin or Terazosin microinjection into bilateral NA, compared to vehicle control injection. (**r**) A diagram illustrating the complete allergen neuroimmune circuit. Scale bars, 100 µm in i, j, m, n, 200 µm in d, f, h. Two-way ANOVA analysis for p and q were performed separately at each MCh concentration. Error bars represent means ± SEM. *p<0.05, ****p<0.0001.

The addition of snRNA-seq signature of this downstream node of the allergen circuit allowed us to define an in silico molecular pathway that starts from the lung and back to the lung (Extended Data Fig. 23a-c). Given our finding that allergen-induced *Fos* activation is reduced in mast cell mutants and following anti-IL-4 antibody neutralization (Fig. 1d-h), we started with key immune signals from type 2 immune cells. Our previous findings also indicate that pulmonary neuroendocrine cells (PNECs) are essential for amplifying allergen-induced responses ^25^. Thus, we added key PNEC products. In the next node the vagal ganglia, we focused on *Trpv1^+^*population because its ablation led to reduced airway hyperreactivity in allergen-challenged mice ^3^. By analyzing published snRNA-seq datasets of the vagal ganglia ^7,8,47,48^, we found multiple corresponding receptors for immune cell and PNEC ligands in *Trpv1^+^*vagal populations (Extended Data Fig. 23a).

We then paired ligands from *Trpv1^+^* vagal neurons and receptors in *Dbh^+^*neurons in the nTS from our snRNA-seq dataset, and ligands from *Dbh^+^*nTS neurons and receptors in the next downstream node using our snRNA-seq datasets (Extended Data Fig. 23b, c). Given that both NA neurons and their downstream airway smooth muscle-innervating postganglionic neurons are *Chat^+^*, the ligands from both of these neuron groups are likely acetylcholine ^49–51^. Taken together, this analysis revealed candidate signals for this neuroimmune circuit that relays from lung immune cells, neurons in the vagal ganglia, nTS and NA of the brainstem, trachea/bronchi, and airway smooth muscle cells.

## Blocking norepinephrine receptors in the NA blunted allergen-induced airway hyperreactivity

To test the function of signals predicted from our ligand-receptor analysis, we focused on the nTS-NA relay. *Dbh* encodes a key enzyme that converts dopamine to norepinephrine, leading to the hypothesis that within the allergen circuit, norepinephrine may be the signal between *Dbh^+^* nTS and NA. From analysis of the NA snRNA-seq dataset and RNAscope assays, among all norepinephrine receptor genes, only *Adra1a* and *Adra1b* are expressed in *Chat^+^* neurons in the NA (Fig. 6i-l). Furthermore, we found that retrograde tracing by tracheal injection of CTB488 labeled *Adra1a^+^* and *Adra1b^+^* neurons in the NA (Fig. 6m, n). These data support that norepinephrine receptor-expressing NA neurons are in the airway circuit, and may receive norepinephrine signals from *Dbh*^+^ nTS neurons in response to allergen challenges.

To test this hypothesis, we addressed whether blocking norepinephrine signal reception in the NA can blunt airway hyperreactivity. Either Prazosin or Terazosin, two norepinephrine receptor antagonists specific for ADRA1, were individually infused into NA bilaterally through implanted cannula (Fig. 6o, Extended Data Fig. 24a). Compared to vehicle control, targeted administration of Prazosin or Terazosin significantly blunted airway hyperreactivity response (Fig. 6p, q, N=6 for HDM+vehicle, N=7 for HDM+Prazosin, N=6 for HDM+Terazosin). These results indicate that the activity of norepinephrine receptor in the NA is necessary for mediating allergen-induced airway hyperreactivity, demonstrating the molecular nature of this key relay in the allergen response circuit.

## Discussion

In this study, our findings delineate a full neural circuit where chronic allergen challenges to lung are transmitted through immune cells such as IL-4 producing immune cells and ascending vagal afferents to a *Dbh^+^* subset of the nTS central integrator, which acts through descending efferent neurons (*Adra1a^+^/Adra1b^+^*) in the NA before projecting back to postganglionic neurons innervating the airway to control airway hyperreactivity (Fig. 6r). This is a circuit identified in wild-type mice following repeated conditioning by allergen, suggesting the presence of an intrinsic molecular and cellular response machinery. Recurring irritation is a common cause of chronic diseases and repeated allergen exposure is central to asthma pathogenesis. Our findings reveal a disease-relevant circuit with players from the immune system, the nervous system and functional cells of the organ.

We found that only repeated, but not single, allergen challenges led to a statistically significant increase in activated nTS neurons. This exemplifies a cardinal feature of chronic conditioning, a behavior that is distinct from mechanisms that drive acute airway constriction ^2,52,53^. These results provide an example of experience-driven neural plasticity in peripheral interoceptive circuits, which are analogous to features classically showcased by central nervous system neurons in learning and memory.

While methacholine can induce acute airway constriction in both groups, repeated allergen challenge led to exacerbated response above the level in naïve animals. Our data demonstrate that this elevated airway resistance was blunted by ablation or chemogenetic inactivation of *Dbh^+^* nTS neurons, demonstrating for the first time the critical role of the nTS neurons in this response. Despite differences, airway hyperreactivity and acute airway constriction may share overlapping nodes. Activation of nervous system in naïve animals, by either electrical stimulation of vagal C-fiber, chemical induction of *MrgprC11^+^* vagal neurons, or optogenetic stimulation of thoracic cholinergic nerves, induced acute airway constriction in the absence of exogenous methacholine ^26,52,53^. These findings suggest that repeated allergen exposures may hijack these existing nodes, condition them through molecular changes or anatomical remodeling, to elevate airway constriction above the threshold level of acute response, mimicking asthma. Our results further define the identity of the brainstem neurons that connect the afferent and efferent nodes to control airway constriction.

While the combined connectivity is necessary and sufficient for allergen-induced hyperactivity, individual nodes of this allergen circuit may also function in other physiological or pathological settings. For example, *Dbh*-expressing nTS neurons were implicated in appetite control by projecting to the lateral parabrachial nucleus or in the arcuate nucleus of the hypothalamus ^54–56^. Dissecting the anatomical connectivity and the pairing of neurotransmitter/neuropeptide to their receptors are key to unraveling the different roles of the same neuron type in distinct circuits. Towards this goal, our snRNA-seq datasets of the nTS and NA provide the basis for dissecting molecular connectivity and specificity. In the context of the allergen circuit, these two datasets, together with existing lung and vagal neuron single cell datasets, and the knowledge that acetylcholine is the likely neurotransmitter from NA to airway smooth muscle cells, complete the connectome of the allergen circuit (Extended Data Fig. 23a-c) ^7,8,47,48^. Two recent studies delineated diverse functions of NA neurons in targeting the esophagus, the heart and/or the lung ^45,46^. It is likely that the cardiopulmonary subset of the NA neurons is responsible for mediating allergen response.

The molecular signatures of the allergen hyperreactivity circuit enables a novel dimension for modulating airway tone in asthma. Neuromodulation could bypass the systemic side effects of current asthma treatments that relay on administration of steroids and biologics. This adaptive circuit showcases the importance of neuroimmune control of organ function. Beyond the lung, mapping how organs act through their peripheral and central nerve connections to receive, process and respond to chronic inputs will open a new dimension in the modulation of disease-relevant interoceptive activities.

## Methods

### Mice

All experimental procedures were performed in the American Association for Accreditation of Laboratory Animal Care (AAALAC)-certified laboratory animal facility at the University of California, San Diego (UCSD). All animal husbandry and experiments were conducted under approved Institutional Animal Care and Use Committee (IACUC) guidelines. Adult (8-16 weeks) mice from strains: C57BL/6J (wild-type, JAX 000664), *c-Kit^w-sh/w-sh^*(JAX 030764)*, Fos^2A-iCreER^* (*TRAP2*, JAX 030323), *Rosa-lxl-tdTomato* (*Ai14*, JAX 007914), *Rosa-lxl-DTR* (JAX 016603), *Chat-cre* (JAX 031661) and *CAG-Sun1/sfGFP* (JAX 030952) lines were purchased from the Jackson lab. *Dbh-cre* (MMRRC 036778) line was purchased from the Mutant Mouse Resource and Research Center (MMRRC). All the *cre* lines we used in this study were kept in B6 background. All *cre* driver lines used are viable and fertile, and no abnormal phenotypes were detected. We crossed *cre* males to B6 wild-type females, and both male and female mice were used in the experiment. Mice were at least six-week-old when performing HDM challenge or doing stereotaxic injection or surgery.

### House Dust Mite (HDM) challenge

Adult mice were anesthetized using isoflurane, 50 μg/20 μl of HDM extract (Dermatophagoides pteronyssinus, GREER Labs) was introduced intranasally weekly continuous for 4 weeks on Days 0, 7, 14, and 21. For controls, 20 μl saline was administered intranasally on the same schedule instead of HDM. Mice were sacrificed 1.5 hours after the 4^th^ challenge for *Fos in situ* hybridization or nTS single-nucleus RNA sequencing, 2 hours after the 4^th^ challenge for FOS antibody immunostaining, 24 hours after the 4^th^ challenge for flexiVent assay, periodic acid–schiff (PAS) staining and quantitative PCR (qPCR), 3 days after the 4^th^ challenge for flow cytometry.

### Histology preparation and PAS staining

Mice were euthanized by CO_2_ inhalation. Lungs were inflated with 4 % paraformaldehyde (PFA) at 35 cm H_2_O airway pressure, fixed overnight and then prepared for paraffin sections at 6 μm. Goblet cells were stained using a PAS staining kit (Sigma).

### Tissue collection and immunofluorescence staining

Mice were euthanized by CO_2_ inhalation. Mice were then transcardinally perfused with Phosphate-buffered saline (PBS) followed by perfusion fixation with 4% PFA. The brainstems were harvested and post-fixed overnight at 4°C in 4 % PFA. After fixing, tissues were washed three times in PBS followed by overnight sucrose dehydration. The brainstem or the lung were then cryo-embedded in OCT compound following standard procedure. Brainstem blocks were sectioned at 20 μm or 40 μm in rostral to caudal sequence, lung blocks were sectioned at 300 μm for parasympathetic neurons/airway staining. All the sections were then processed for immunostaining following a standard protocol. Primary antibodies used include rabbit anti-c-FOS (SYSY, 226 003), rabbit anti-DBH (Sigma, AB1585), rabbit anti-Dsred (Takara, 632496), rabbit anti-VAChT (SYSY, 139 103), mouse anti-alpha Smooth Muscle Actin-FITC (Sigma, F3777). Secondary antibodies used include goat anti-rabbit FITC, goat anti-rabbit Cy3, goat anti-rabbit Cy5 (all from Jackson Immuno Research Labs). Slides were mounted using Vectashield (Vector Labs) and imaged using the Olympus VS200 Slide Scanner or the Leica SP8 confocal microscope. For quantification of FOS^+^ (Fig. 1c, e, h, j, Extended Data Fig. 3c, f, i, Extended Data Fig. 13c) or tdTomato^+^ (Fig. 1p and Extended Data Fig. 5c) cells in the nTS, or *Fos^+^* cells after RNAscope *in situ* hybridization (Extended Data Fig. 12f), we did serial section of the whole nTS and counted from three cryosections (three sections were chosen coordinate between Bregma -7.20 mm and -8.08 mm with the same stereotaxic, where HDM-induced FOS^+^ cells are elevated) with the maximal numbers of FOS^+^ cells and count the average number as one data point for one animal. We used Qupath for overlap quantification in the nTS (Fig. 3d, f).

### RNAscope *in situ* hybridization

To prepare nTS sections for RNAscope *in situ* hybridization, fresh brainstems were cryo-embedded in OCT compound. Sections at 12 μm were prepared using a cryostat, collected on Superfrost Plus slides, and left to dry in the cryostat for at least 30 minutes before staining. All staining procedures were performed using RNAscope Fluorescent Multiplex Kit (Advanced Cell Diagnostics, 320850) following the manufacturer’s instruction. Briefly, sections were post-fixed in 4% PFA for 1 hour at room temperature, washed in PBS, dehydrated in a series of ethanol washes, and then dried. A hydrophobic barrier was drawn around the section with an ImmEdge pen (Vector Lab, H-4000). The sections were treated with Protease IV in a HybEZ Humidity Control Tray for 30 minutes at room temperature, incubated with target probes in a HybEZ Oven for 2 hours at 40°C, and then treated with Hybridize Amp buffer. Slides were mounted using Fluoromount-G with DAPI (Southern Biotech). The following probes from Advanced Cell Diagnostics were used: Mm-*Fos* (316921), Mm-*Dbh* (407851), Mm-*Gil3* (445511), Mm-*Sncg* (482741), Mm-*Slc17a7* (431201), Mm-*Mafa* (556931), Mm-*Fos* (316921), Mm-*Cnr1*(420721), Mm-*Th* (317621), Mm-*Cck* (402271), Mm-*tdTomato* (317041), Mm-*Chat* (408731), Mm-*Adra1a* (408611), Mm-*Adra1b* (413561), Mm-*Adra2a* (425341), Mm-*Adra2b* (425321), Mm-*Adrb1* (449761), Mm-*Adrb2* (449771) and Mm-*Adrb3* (495521).

### Vagotomy

Mice were anesthetized with a mixture of ketamine (100 mg/kg) and xylazine (10 mg/kg) via intraperitoneal injection. Fully anesthetized mice were secured in the sternal position to the homoeothermic blanket using lab tape. Spread 75 % EtOH on the throat of the mouse to wet the fur. Lift the skin to make a vertical cut (1 cm) on the throat, isolate one side of vagal nerve and tease away from carotid artery using small, curved forceps. Pass fine, curved forceps underneath the vagal nerve and slowly open forceps to separate the nerve fiber from the surrounding tissue. Unilateral vagotomy was conducted by lifting up the vagal nerve and cutting with straight scissors. Control sham operations are simulated by lifting the vagal nerve and releasing it intact. After vagotomy, the wounds were closed carefully with sterilized dissecting scissors and suture strings. The area will be cleaned with 75% EtOH and disinfected with Povidone-Iodine. The time between fully anesthesia and completion of the surgery should take less than 10 minutes.

Buprenorphine SR (0.1 mg/kg, subcutaneous injection) was used for pain relief following surgery. One week after surgery, mice were used for allergen challenge.

### 4-OH tamoxifen injection and *TRAP2* labeling

4-hydroxytamoxifen (4-OHT, Sigma H6278) was dissolved at 20 mg/mL in ethanol by shaking at 37°C for 15 minutes and was then aliquoted and stored at -20°C for up to several weeks. Before use, 4-OHT was redissolved in ethanol by shaking at 37°C for 15 minutes, a 1:4 mixture of castor oil: sunflower seed oil (Sigma, 259853 and S5007) was added to give a final concentration of 10 mg/mL 4-OHT, and the ethanol was evaporated by vacuum under centrifugation. All injections were delivered intraperitoneally to adult *TRAP2* mice. Mice were kept in their home cage for additional 1 week to allow the expression of tdTomato before treatment.

### CLARITY-based brain clearing

A hydrogel based on 1 % acrylamide (1 % acrylamide, 0.125 % Bis, 4 % PFA, 0.025 % (w/v) VA-044 initiator, in 1X PBS) was used for all CLARITY preparations. Mice were transcardially perfused with 4 % PFA. After perfusion, brains were post-fixed in 4% PFA overnight at 4°C and then transferred to 1 % hydrogel for 48 hours to allow monomer diffusion. The samples were degassed and polymerized (4-5 hours at 37°C) in a 50 ml tube. The brains were removed from hydrogel and washed with 200 mM NaOH-Boric buffer (pH=8.5) containing 8 % sodium dodecyl sulfate (SDS) for 6-12 hours to remove residual PFA and monomers. Brains could then be transferred to a flow-assisted clearing device using a temperature-control circulator or a simpler combination of 50 ml tube and heated stirring plate. 100 mM Tris-Boric Buffer (pH=8.5) containing 8 % SDS was used to accelerate the clearing (at 40°C). A whole mouse brain can be cleared in 12-16 days. After clearing, the brain was washed in PBST (PBS+0.2% Triton X) for at least 24 hours at 37°C to remove residual SDS. Brains were incubated in a refractive index matching solution (RI=1.45) for 8 hours (up to 1 day) at 37°C and then 6-8 hours at room temperature. After the incubation, brainstem hemispheres were imaged using confocal microscope.

### Brainstem stereotaxic injection

Fully anesthetized mice (ketamine at 100 mg/kg and xylazine at 10 mg/kg via intraperitoneal injection) were placed in a stereotaxic frame with the head angled at 45° and body temperature maintained with a heating pad. A midline incision was made though the animal’s skin, posterior neck muscles, and dura mater to expose the medulla between the occipital bone and C1 vertebra. Based on the stereotaxic coordinates of mouse brain ^57^ and using Obex as a reference point, bilateral injections were made into the nTS (0.1 mm rostral to Obex, 0.2 mm lateral to midline, 0.25 mm under the medullary surface), the DMV (0.05 mm caudal to Obex, 0.1 mm lateral to midline, 0.1 mm under the medullary surface), or the NA (0.65 mm caudal to Obex, 1.25 mm lateral to midline, 0.45 mm under the medullary surface) using a calibrated glass micropipette attached to a Nanoject II injector (Drummond) microprocessor pump (Pneumatic PicoPump; WPI). Over 5 minutes, DTX (2 ng/200 nl), anti-DBH-SAP (42 ng/200 nL, advanced targeting system, IT-03), blank-SAP (advanced targeting system, IT-21), or viruses (50 nl for nTS or DMV, 20 nl for NA, Boston Children’s Hospital Vector Core) were gradually injected at the targeted site of *TRAP2* mice, wild-type mice, *Dbh-cre* or *Chat-cre* mice, respectively. Viruses include AAV2/9-flex-tdTomato (1.3x10^13^ genome copies (gc)/ml), AAV2/8-flex-hM4D(Gi)-mCherry (2.07x10^13^ gc/ml), AAV2/8-DIO-hM3D(Gq)-mCherry (8x10^12^ gc/ml). After injection, the glass micropipette was left in place for 10 minutes to limit nonspecific spread of the toxin or virus. The micropipette was pulled out very slowly. Buprenorphine SR (0.1 mg/kg, subcutaneous injection) was used for pain relief following surgery. The mice were allowed to recover from anesthesia on a heating blanket before returning to the home cage.

### Airway hyperreactivity assayed by flexiVent

Anesthetized (100 mg/kg ketamine and 10 mg/kg xylazine, intraperitoneal injection) mice were paralyzed with acepromazine (10 mg/kg, intraperitoneal injection) and then tracheotomized with a 20G sterile catheter, attached to a flexiVent pulmonary mechanics apparatus (SCIREQ). Mice were ventilated at a tidal volume of 9 ml/kg with a frequency of 150 bpm. Positive end-expiratory pressure was set at 300 mm H_2_O. Lung resistance (Rrs) and elastance (Ers) of the respiratory system were determined in response to aerosolized methacholine challenges (0, 6, 12 and 24 mg/ml). All mice used were adults of similar age and weight. The weight of each animal was entered into flexiVent at the start of each round of assay. As part of the flexiVent program, pre-scans were carried out that led to calculation of lung size. Methacholine was dissolved in sterile saline. The nebulizer was on for 10 seconds to deliver each dose of methacholine. The mean maximal elastance and resistance of twelve measurements by doses was calculated.

Samples were compared using two-way ANOVA. Statistical analysis at each methacholine concentration was performed separately. Error bars represent means ± SEM. Statistical significance called at p<0.05.

### Tissue dissociation and flow cytometry

Three days after last challenge, mice were anesthetized (100 mg/kg ketamine and 10 mg/kg xylazine, intraperitoneal injection) and then injected with AF700-counjugated CD45 through intravenous injection to distinguish circulating immune cells and immune cells resident within the lung. Mice were euthanized 5 minutes later for lung harvest. Whole lungs were mechanically dissociated in GentleMACS C tubes (Miltenyi Biotec) containing 5 ml of PRMI 1640 (Thermo Scientific) with 10 % FBS, 1 mM HEPES (Life Technology), 1 mM MgCl_2_ (Life Technology), 1 mM CaCl_2_ (Sigma-Aldrich), 0.525 mg/ml collagenase/dispase (Roche) and 0.25 mg DNase I (Roche) by running mouse lung 1-2 program on GentleMACS (Miltenyi Biotec). Lung pieces were then digested by shaking (∼150 rpm) for 30 minutes at 37°C. After incubation, Lung pieces were mechanically dissociated further using mouse lung 2-1 program on GentleMACS, followed by straining through a 70 mm filter. Red blood cells were removed by adding 1 mL RBC lysis buffer (Biolegend) to each tube and incubate at room temperature for 1 minute.

The single-cell suspensions from above were then pelleted (1500 rpm, 4°C, 5 minutes), counted with hemocytometer and diluted to around 1 × 10^6^ cells per ml. They were stained with Fc blocking antibody (5 mg/ml, BD) at 4°C for 30 minutes. The cells were washed with DPBS and then incubated with surface marker antibody cocktail for 30 minutes at 4°C. For flow cytometry of lung myeloid, the following antibodies were used (all antibodies were from BioLegend): 1:100 AF488-conjugated anti-F4/80; 1:500 BV510-conjugated anti-CD45; 1:500 BV605-counjugated anti-Siglec-F; 1:100 V450-counjugateed anti-TCR-beta; 1:200 V450-counjugated anti-B220; 1:1000 PerCP-Cy5.5-conjugated anti-Ly6C; 1:2000 PE-conjugated anti-MHC-II; 1:1000 APC-conjugated anti-CD11c; 1:1000 PE-Cy7 conjugated anti-Ly6G; 1:2000 PE-CF594-counjugated CD11b. For flow cytometry of lung lymphoid, the following antibodies were used (all antibodies were from BioLegend): 1:200 FITC-conjugated anti-CD45; 1:100 AP-Cy7-counjugateed anti-IL7Ra; V450-counjugated Lineage mix (1:200 anti-CD19; 1:500 anti-CD11c; 1:500 anti F4/80; 1:100 anti-NK1.1; 1:100 anti-TER119; 1:100 anti-TCR gamma delta); 1:100 BV510-conjugated anti-ST2; 1:200 PE-Cy7-counjugated anti-TCR-beta; 1:100 BV604-counjugated anti-CD4; 1:2000 PerCP-Cy5.5-conjugated anti-CD90.2; 1:100 APC-conjugated anti-cKit; 1:2000 PE-CF594-counjugated anti-CD11b; 1:200 PE-conjugated anti-MAR.1. The cells were then stained using live/dead dye (1:1000, Ghost Dye Red 780, TONBO biosciences for myeloid, 1:500, Ghost Dye Violet 450, TONBO biosciences for lymphoid) for 30 minutes at 4°C. The cells were fixed using BD Stabilizing Fixative for 30 minutes at 4°C before transferring to FACs tubes. Flow cytometry was analyzed on BD FACS Canto RUO - ORANGE with three lasers (405 nm, 488 nm and 640 nm) using the Flow Cytometry Core at the VA San Diego Health Care System and the San Diego Veterans Medical Research Foundation. All data were further analyzed and plotted with FlowJo software (Tree Star). Eosinophils, innate lymphoid cells (ILC2s) and Th2 cells were gated on live, resident CD45^+^ singlets.

### Isolation of nuclei from frozen nTS

We dissected out the whole nTS region from adult C57BL6/J male mice brainstem. Mice were euthanized using CO_2_ inhalation. Brains were acutely harvested from two groups of wild-type males (WT), or from two adult saline-or HDM-challenged males at 1.5 hours after the 4^th^ challenge, respectively (two technical replicates in every group). The nTS was visualized by microscopy and harvested based on anatomical landmarks. Dissected nTS were put in liquid N_2_ immediately and store at -80°C or proceed directly to nuclei isolation. On the day of nuclei dissociation, plunge the dounce and pestle in ice. Transfer douncing buffer (DB, 0.25 M sucrose, 25 mM KCl, 5 mM MgCl_2_ and 10 mM Tris-HCl, pH=7.5) to the dounce in ice and let them chill. Transfer dissected nTS into dounce which includes 1 mL DB with 1 mM DTT (D9779, Sigma), 1x cOmplete EDTA-free protease inhibitor (05056489001, Roche, DB-DP) with 0.1% Triton-X. Use loose pestle 5-10 times gently followed by 15-20 stokes with tight pestle, without introducing bubbles. After filter with 30 μm CellTrics, transfer the contents to pre-chilled low bind Eppendorf tube. Spin at 1,000 g for 10 minutes at 4°C. Remove supernatant and gently resuspend in 1 mL of DB-DP. Spin at 1,000 g for 10 minutes at 4°C. Remove supernatant and gently resuspend in 1 mL permeabilization buffer (BSA, 5% IGEPAL-CA630 (I8896, Sigma), 0.2% DTT, 1 mM cOmplete EDTA-free protease inhibitor, 1xPBS). Resuspend in 50-500 µL of 1.11x tagmentation buffer (66 mM Tris-acetate (pH=7.8, BP-152, Thermo Fisher Scientific), 132 mM K-acetate (P5708, Sigma), 20 mM Mg-acetate (M2545, Sigma) and 32 mM DMF (DX1730, EMD Millipore). Count with hemocytometer.

### Single-nucleus RNA sequencing and data analysis

Single-nucleus RNA sequencing experiments were provided by the Center for Epigenomics, University of California, San Diego. nTS nuclei were processed into cDNA libraries using Chromium Single Cell 3’ v3 kit (10XGenomics). Sequencing was carried out on NovaSeq (Illumina) platform at the Institute for Genomic Medicine, University of California, San Diego (UCSD). CellRanger package (version 3.0.2) was used to align the raw reads onto the mouse reference genome (GRCm38) and generate the feature-barcode matrix. CellBender was used next to model and remove systematic biases and background noise, and to produce improved estimates of gene expression. Next, R package Seurat (version 4.0) was used to perform data quality control, normalization, principal components analysis and uniform manifold approximation and projection (UMAP method). For raw data from WT, Saline or HDM conditions, we firstly used SCTransform to regress our batch effects and did doublets removal separately. We then integrated all the data from three conditions (WT, Saline, HDM) using Harmony. Cells with 3,500 to 5,500 unique feature counts were considered high quality cells and used for further analysis. The integrated dataset was normalized, scaled, and then analyzed for variable genes using Seurat’s NormalizeData, ScaleData, and FindVariableFeatures functions, respectively. A total of 2,000 top variable features were identified by function FindVariableFeatures and selected for subsequent principal components analysis. Top 20 significant components were chosen to conduct cell clustering by using the algorithm of uniform manifold approximation and projection (UMAP) with default settings. The expression level and feature of selected genes were profiled and visualize by R package ggplot2 (Version 3.3.2). We then read snRNA-seq data from WT condition and profiled dot plot and UMAP plot of marker gene expression in all the nTS clusters (Fig. 2a-q, Extended Data Fig. 7a-o, Extended Data Fig. 8a). We compared snRNA-seq data from Saline and HDM conditions and profiled dot plot of *Fos* in all the nTS clusters (Fig. 3a, b, Extended Data Fig. 8b).

### Quantitative PCR (qPCR)

Total RNA was extracted from lungs using Trizol (Invitrogen) and RNeasy Mini RNA extraction kit (Qiagen). RT-PCR was then performed to obtain corresponding cDNA using iScript Select cDNA Synthesis Kit (Bio-Rad). qPCR was performed by CFX ConnectTM system (Bio-Rad) using SYBR Green (Bio-Rad). At least three technical and three biological replicates were performed for each gene if not otherwise notated. Samples were compared using unpaired student’s t-test. Error bars represent means ± SEM. Statistical significance called at p<0.05.

Primers

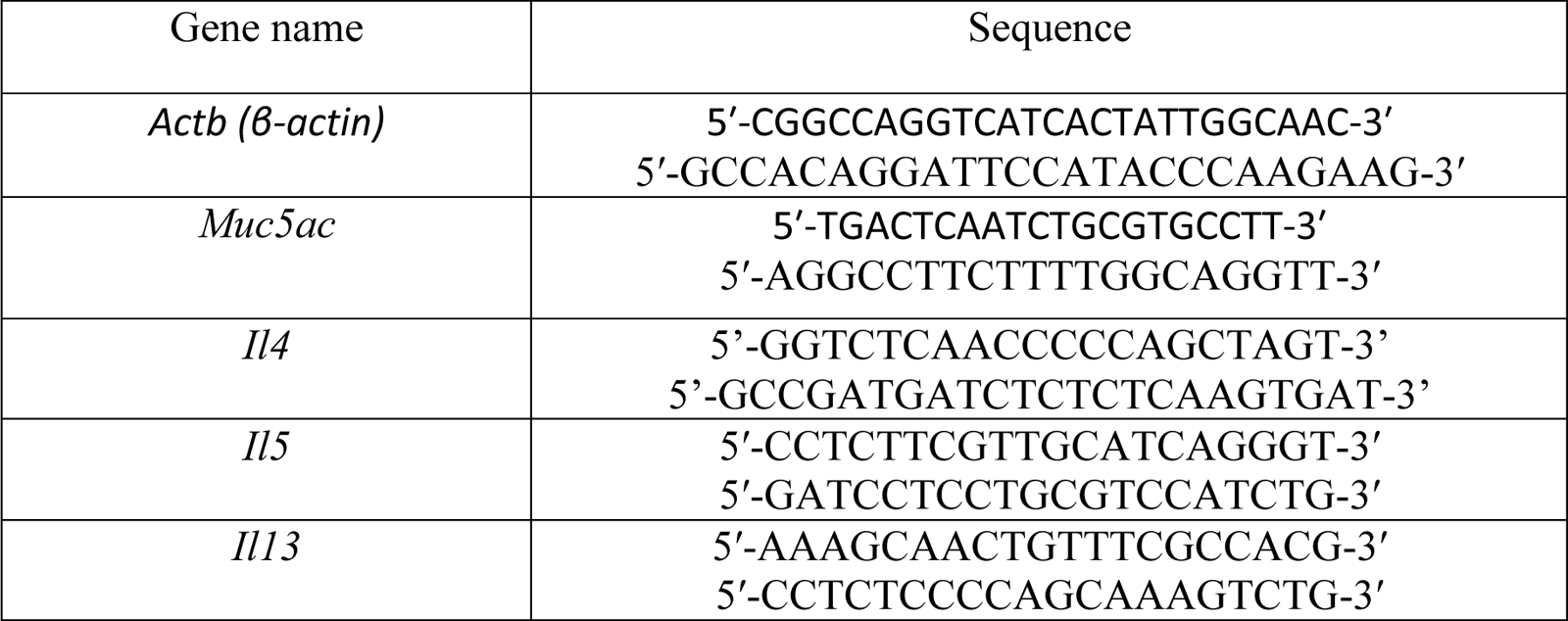

### nTS-targeted SAP injections

Mice received bilateral injections of either unconjugated blank-SAP or SAP conjugated to an antibody directed to DBH (DBH-SAP; 42 ng/200 nL, pH=7.4; MAB394; Millipore, Temecula, CA, USA), using a Hamilton syringe directed toward the nTS (0.1 mm rostral to Obex, 0.2 mm lateral to midline, 0.25 mm under the medullary surface). Blank-SAP cannot enter cells and is non-toxic to neurons, serving as the appropriate control for DBH-SAP administration. Over 5 minutes, 50 nL of blank-SAP or DBH-SAP was gradually injected to the nTS. DBH-SAP was previously validated to specifically ablate DBH^+^ neurons with no effects on neighboring neurons^41^. Upon specific binding to DBH at the synaptic cleft of DBH^+^ cells, the complex including conjugated anti-DBH-SAP antibody is endocytosed and transported back to the cell body ^58^. In the cell body, the SAP toxin inactivates ribosomes, leading to cell death ^59^. As previously published studies indicate that 2 weeks is necessary to eliminate *Dbh^+^* neurons using DBH-SAP ^58^, mice were injected 1 week before the 1^st^ HDM sensitization and 2 weeks before the 2^nd^ challenge.

### Plethysmography

Ventilatory parameters during normoxia was measured in unrestrained male mice using a whole-body barometric plethysmograph (500 mL) modified for continuous flow ^60–62^. Flow was maintained constant through the chamber while a pressure transducer (45 mMP with 2 cm H_2_O diaphragm, Validyne, Northridge CA) recorded the changes attributable to the warming and expansion of inhaled gases. On the experimental day, the mice were weighed and sealed into an individual plethysmograph chamber along with a temperature and humidity probe (Thermalert TH5, Physitemp, Clifton NJ). A constant gas flow (335 ml/min) was delivered with a rotameter (603, Matheson, Montgomeryville PA) and measured with a flow meter (Sables System International, Inc. Las Vegas, NV) upstream of the chamber. Gases exited the chamber through a valve and into a vacuum pump (Model 25, Precision Scientific Co, Chicago IL) to isolate pressure changes from breathing in the chamber during constant flow with high input and output impedances. This also allows us to maintain chamber pressure near-atmospheric pressure and reference pressure measurements (Validyne) in the chamber to atmosphere.

All ventilatory parameters were recorded on an analog-digital acquisition system (PowerLab 8SP, AD Instruments) and analyzed with the LabChart 8-Pro Software, sampling at a rate of 1 kHz. Mice were allowed to acclimatize to the chamber and constant air flow for 45 minutes (air with 21% of O_2_) and then exposed to a 5-minute challenge of hypoxic gas (10% O_2_) to test responsiveness. After that we exposed the mice to normoxic gas for 15 minutes. We analyzed a minimum of 30 seconds as part of the region of interest between 10 and 15 minutes of the las exposure to normoxic gas. The frequency (breaths/min) and tidal volume (ml) were calculated from cyclic peaks in the plethysmograph pressure pulses, and tidal volume was measured using ml calibration pulses and the equations were developed by Drorbaugh and Fenn ^63^. We measured minute ventilation (VI), respiratory frequency (Rf) and tidal volume (Vt). All of them were measured under 21% of oxygen. Minute ventilation (VI) is the product of frequency and tidal volume, which was normalized to body mass ventilation (ml/min·kg). We also calculated oxygen consumption (V_O2_) and carbon dioxide production (V_CO2_) during the ventilation measurement by recording the inspired and expired oxygen and carbon dioxide fractions were measured with an O_2_/CO_2_ analyzer (FOXBOX, Sables System International, Inc. Las Vegas, NV) sampling from the chamber, and then calculating inspired and expired O_2_ and CO_2_ fractions, chamber flow rate and correcting for water vapor and respiratory quotient (eqn. 10.6 and 10.7 in J.R.B. Lighton, Measuring Metabolic Rates, Oxford Univ. Press, Oxford, 2008). Incurrent gases are dry from tanks. Water vapors were scrubbed from excurrent gas when it was analyzed with the O_2_/CO_2_ analyzer. Thus, we calculated the ratio of VI / V_O2_ and VI / V_CO2_ to have a more precise estimation of the mice ventilation avoiding confounding factors produced by changes in metabolic rate.

### CUBIC tissue clearing and light sheet fluorescence microscopy

CUBIC (Clear, Unobstructed Brain/Body Imaging Cocktails and Computational analysis protocol) buffers were prepared according to Susaki et al ^64^. After sufficient tissue clearing in R1 buffer, tissues were embedded in 2 % low melting agarose and then incubated in R2 solution at room temperature overnight prior to imaging. Cleared samples were imaged using a Zeiss Z.1 light sheet fluorescence microscope (LSFM). Lung samples were imaged using a 2.5x objective (LSFM 2.5x NA 0.1) and a 1.45 2.5X CLARITY specific chamber (Translucence Biosystems).

### Isolation of nuclei from frozen NA

We dissected out the NA region (GFP^+^) from adult *Chat-cre; CAG-Sun1/sfGFP* mice brainstem under fluorescence microscope. Mice were euthanized using CO2 inhalation. Brains were acutely harvested. The NA was visualized by nucleus localized GFP fluorescence signals and harvested based on anatomical landmarks. We collected GFP+ neurons around NA region and avoided DMV and 12N regions which also express Chat. NA from 7 mice were dissected and pulled together and put in liquid N_2_ immediately and store at -80°C or proceed directly to nuclei isolation. We used the same dounce dissociation method that we isolated nTS nuclei (see method-Isolation of nuclei from frozen nTS).

### NA single-nucleus RNA sequencing and data analysis

NA nuclei were processed, sequenced and analyzed using the same kit, platform and software that were used for nTS (see method-nTS single-nucleus RNA sequencing and data analysis). For raw data, we firstly confirmed the mitochondrial counts was < 5%, and cells above 200 unique feature counts were considered high quality cells and used for further analysis.

### Under trachea injection

Mice were anesthetized with a mixture of ketamine (100 mg/kg) and xylazine (10 mg/kg) via intraperitoneal injection. After fully anesthetized, mice were placed ventral side up in a stereotaxic frame with body temperature maintained with a heating pad. Ophthalmic ointment was applied to maintain eye lubrication. The extra thoracic trachea was exposed via a ventral incision in the animal’s neck. The trachea was lifted up by forceps carefully without bleed the trachea. Using a 10 μl Hamilton glass micropipette fitted with a 32G needle, 10 μl CTB488 (concentration at 1 mg/ml, Thermo Scientific, C22841) was injected into the dorsal side of the trachea from bilateral sides, 3 to 5 cartilage rings caudal to the larynx. The injection was done within 15 minutes per mouse. Mouse skin was sutured and recovered on thermal pad before returned to housing. Buprenorphine SR (0.1 mg/kg, subcutaneous injection) was used for pain relief following surgery. For brainstem collection, mice were sacrificed 1 week after CTB488 injection.

### Cannula implant and microinjection into bilateral NA

Mice were anesthetized with a mixture of ketamine (100 mg/kg) and xylazine (10 mg/kg) via intraperitoneal injection. After fully anesthetized, mice were mounted in a stereotaxic frame. A midline incision was made to reveal Bregma, the skull was cleaned with hydrogen peroxide, and small holes were drilled through the skull at the designated stereotaxic coordinates of NA^57^.

Bilateral guide cannula affixed with two 26G stainless-steel tubing (P1 Technologies Inc., Roanoke, VA, USA) was stereotaxically implanted 0.5 mm above the NA region (0.7 mm caudal to Bregma, 1.25 mm lateral to midline, 0.4 mm under the dura surface). Guide cannulas were secured to the skull using super glue and dental cement. Matching dummy cannula were inserted into the guide cannula and secured with a dust cap to ensure guide cannula patency. Post-operative analgesia was provided with Buprenorphine SR (0.1 mg/kg, subcutaneous injection). Mice recovered in their home cages for one week before challenge and drug delivery.

Starting from the baseline measurement of flexiVent assay, Prazosin (0.4 mg/ml in ultra-pure water, P7791, Sigma), or Terazosin (0.4mg/ml in ultra-pure water, T4680, Sigma) or ultra-pure water control was continuously microinjected (total volume 1 μl, continuously injecting for 17.5 minutes) into bilateral NA through the guided cannula using two-channel syringe pump (R462, RWD, USA). For all the mice for NA microinjection, we kept the brainstem and confirmed targeted cannula implantation in NA region through cryosection.

### Statistical analysis

All statistics were calculated using Microsoft Excel and performed using GraphPad Prism (GraphPad Software Inc., CA, USA). Unpaired student’s t-test, one-way or two-way ANOVA Tukey’s multiple comparisons test were used as indicated in figure legends to examine significant differences across mice and experimental conditions. Values of p<0.05 were deem significant and lack of significant differences (p>0.05) are indicated by N.S. (not significant).

## Data and materials availability

The accession number for the raw and fully processed nTS single-nucleus RNA sequencing data reported in this paper is GEO: GSE200003. The accession number for the raw and fully processed NA single-nucleus RNA sequencing data reported in this paper is GEO: GSE211538. Both are publicly available upon publication.

## Acknowledgments

We thank members of the Sun lab for inputs and discussions. We thank Drs. Takaki Komiyama, David Kleinfeld and Byungkook Lim at the University of California San Diego (UCSD), Dr. Samuel Pfaff at the Salk Institute for inputs and discussions. We thank the Center for Epigenomics at UCSD for performing the single nucleus RNA-seq experiments. Work at the Center for Epigenomics was supported in part by the UCSD School of Medicine. This publication includes data generated at the UCSD Institute for Genomic Medicine utilizing an Illumina NovaSeq 6000 that was purchased with funding from a National Institutes of Health SIG grant (#S10 OD026929). Flow cytometry was performed with the support of the Flow Cytometry Research Core at the San Diego Center for AIDS Research (P30 AI036214), the VA San Diego Health Care System, and the San Diego Veterans Medical Research Foundation. We thank Jennifer Santini and Marcella Erb at the UCSD School of Medicine Microscopy Core (National Institute of Neurological Disorders and Stroke Grant P30 NS047101) for assistance with imaging. Fundings were provided by the National Institutes of Health 1R01AT011676 (X.S.), National Institutes of Health OT2OD023857 (X.S.), Tobacco-Related Disease Research Program (TRDRP) T29IR0475 (X.S.) and American Heart Association 19POST34450103 (Y.S.).

## Author contributions

Y.S. and X.S. conceived and designed experiments; Y.S. performed experiments; Y.S., A.N. and X.S. interpreted data; Y.S. prepared figures and drafted manuscript. J.X. helped with *TRAP2* and flexiVent experiments, and performed PAS staining; Z.Z., H.Y. and Y.S. analyzed single-nucleus RNA-seq data and generated figures; V.N. and L.Y. performed CLARITY of *TRAP2* mice and did confocal imaging; B.D. designed antibody cocktails for flow cytometry; E.M. performed plethysmography and analyzed the data; Y.S. and X.S. edited and revised the manuscript with input from the other authors; Y.S., J.X., Z.Z., H.Y., V.N., B.D., E.M., L.Y., A.N. and X.S. approved final version of manuscript.

## Materials & Correspondence

Correspondence and requests for materials should be addressed to Xin Sun.

All figures and extended data figures have been deposited in figshare, and the private sharing links are provided here to access.

Fig. 1: https://figshare.com/s/c93fff81efdb06965a55

Fig. 2: https://figshare.com/s/45c42b6036ecf8f24a7d

Fig. 3: https://figshare.com/s/366a0e6b67eb9abef37f

Fig. 4: https://figshare.com/s/37ccb9018f4653d0ecfd

Fig. 5: https://figshare.com/s/95a526de44919eeaff88

Fig. 6: https://figshare.com/s/1bd52dd2ff58a5853cca

Extended Data Fig. 1: https://figshare.com/s/67ca20595b020a9b61cf

Extended Data Fig. 2: https://figshare.com/s/f8a2cf4e4029a0bf806d

Extended Data Fig. 3: https://figshare.com/s/2ce8910e15a77da11a90

Extended Data Fig. 4: https://figshare.com/s/07566d37b1a37b109190

Extended Data Fig. 5: https://figshare.com/s/f0d231f3fd9c71da8167

Extended Data Fig. 6: https://figshare.com/s/be7a508b61b645125db9

Extended Data Fig. 7: https://figshare.com/s/480e03f3d08e98464bd4

Extended Data Fig. 8: https://figshare.com/s/4f1489d96416fc9071d6

Extended Data Fig. 9: https://figshare.com/s/1a1bc40336f6c2f61996

Extended Data Fig. 10: https://figshare.com/s/4fe23454fc61b4e7388a

Extended Data Fig. 11: https://figshare.com/s/612f575fec98429ae61e

Extended Data Fig. 12: https://figshare.com/s/05e0065abc0baac90be1

Extended Data Fig. 13: https://figshare.com/s/2105e42d4cf4c82b2a2d

Extended Data Fig. 14: https://figshare.com/s/dbefeabb2dcfe5e58731

Extended Data Fig. 15: https://figshare.com/s/4e0c530edb25cd77617e

Extended Data Fig. 16: https://figshare.com/s/79b7907b8190d8ce2e7e

Extended Data Fig. 17: https://figshare.com/s/1c1bfd80b34556126476

Extended Data Fig. 18: https://figshare.com/s/70f4f3730099261788be

Extended Data Fig. 19: https://figshare.com/s/1903a369125fcfb9e701

Extended Data Fig. 20: https://figshare.com/s/d50fe7c0874ca27fa01d

Extended Data Fig. 21: https://figshare.com/s/086052bb8b27fe4dd2bb

Extended Data Fig. 22: https://figshare.com/s/a19716980cd22d3cf2cf

Extended Data Fig. 23: https://figshare.com/s/ad9bc446b926845a0942

Extended Data Fig. 24: https://figshare.com/s/789a21c020440c3fdbfb

## Extended Data-figures and figure legends

**Extended Data Fig. 1.**
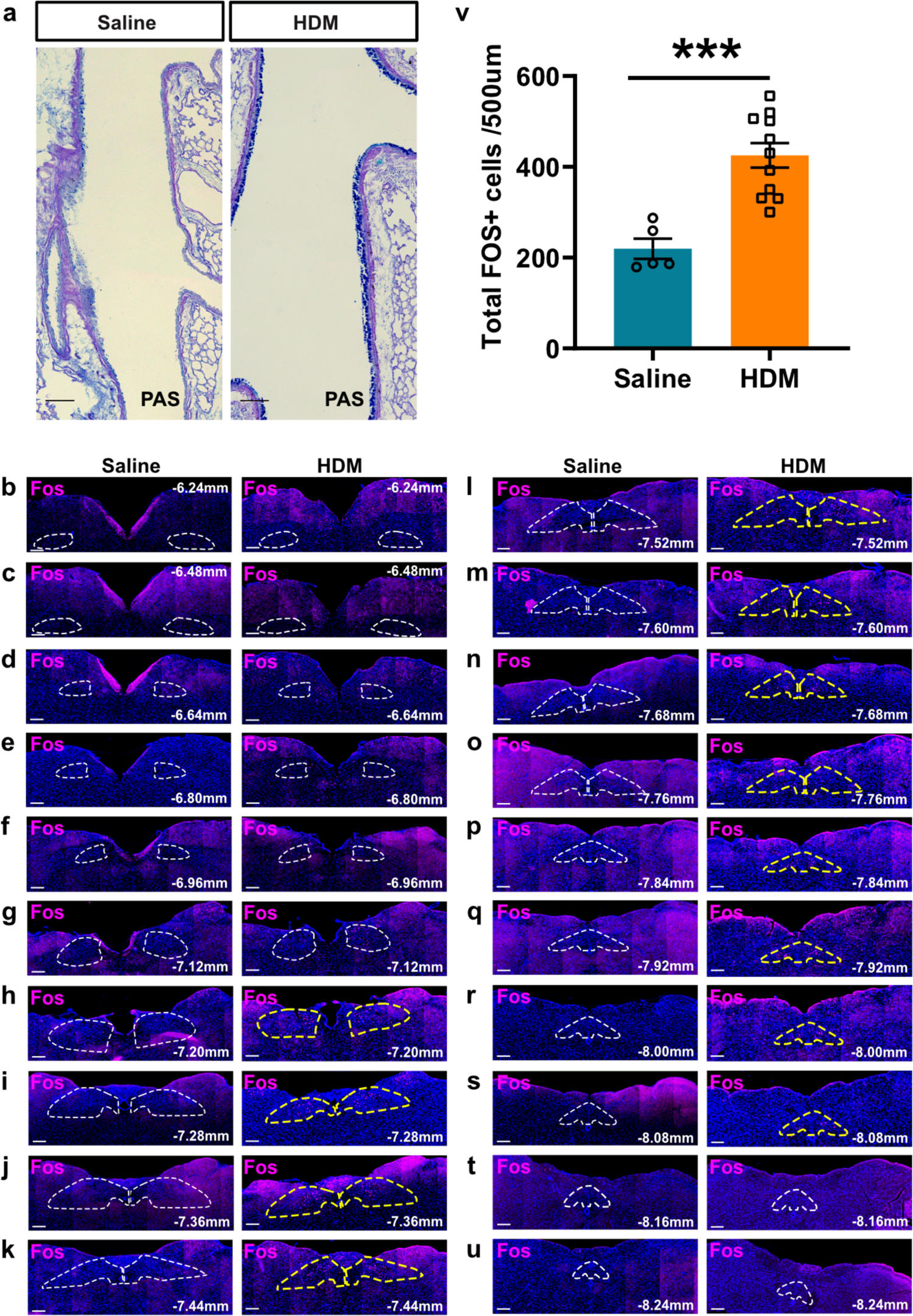
nTS neurons were activated after allergen challenge in lung. (**a**) PAS staining shows that HDM regimen effectively induced mucus-secreting airway goblet cells as expected. (**b-u**) FOS antibody staining two hours after the 4^th^ challenges in serial sections of the whole brainstem. Increased FOS^+^ cells were enriched in the dorsomedial part of the nTS region from Bregma -7.20 mm to -8.08 mm in HDM-treated mice (yellow outlined) compared to the saline-treated control mice. (**t**) Quantification of total number of FOS^+^ cells after HDM or Saline control in the whole nTS regions (Bregma -6.24 mm to -8.24 mm). Each serial section was at 20 µm thick, 80 µm interval in-between sections, so the number represent ¼ (500 µm) of the whole nTS regions. Each data point represents result from one animal. Scale bars, 200 µm in b-u. Unpaired student’s t test was used for t. Error bars represent means ± SEM. ***p<0.001.

**Extended Data Fig. 2.**
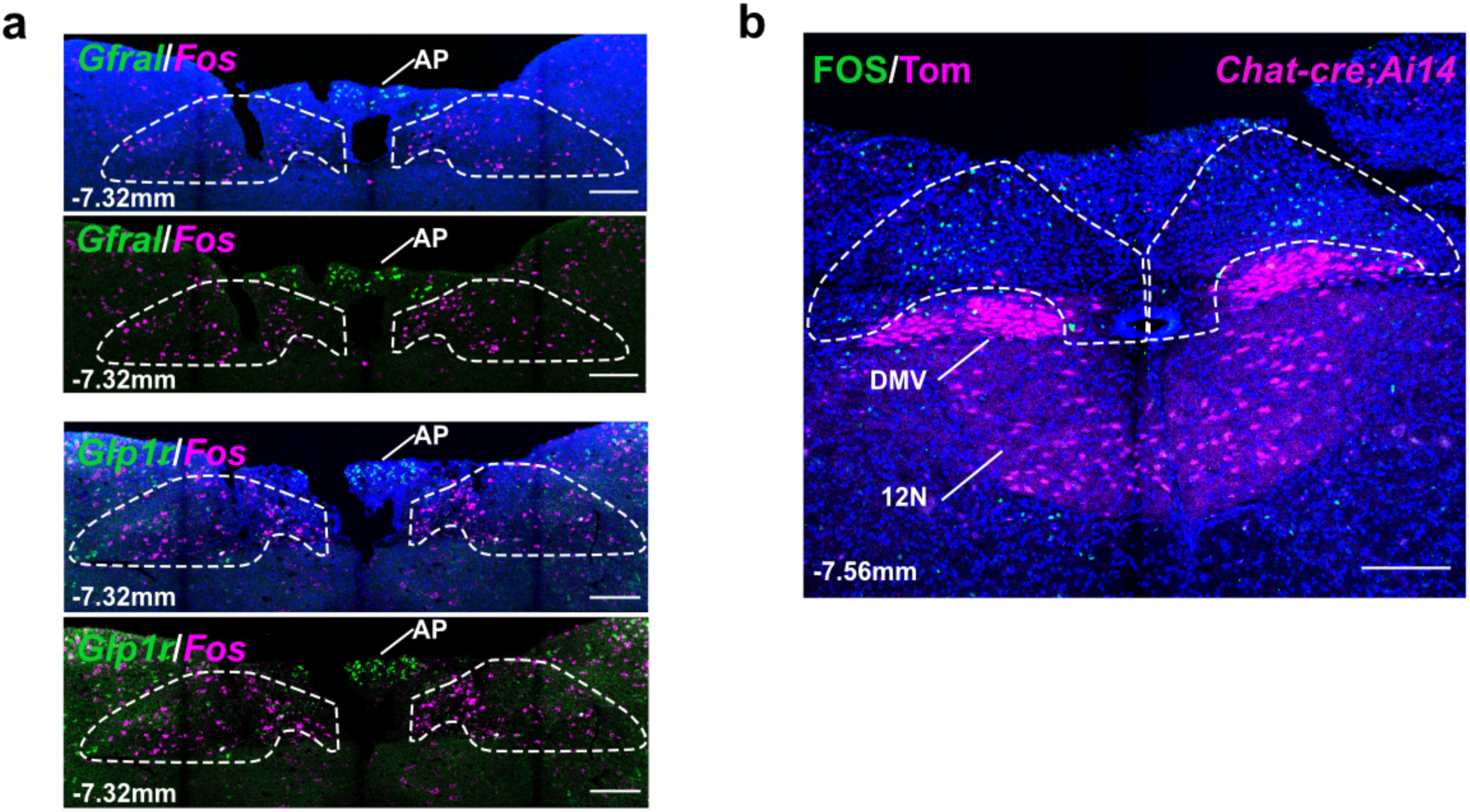
Allergen-activated neurons were not located in the adjoining AP, DMV or 12N regions in the brainstem. (**a**) *Fos in situ* hybridization from brainstem section of naïve mice after HDM challenge in lung showing most *Fos* RNA signals do not overlap with genes (*Gfral* or *Glp1r*) that are enriched in the AP. (**b**) FOS antibody staining from brainstem section of *Chat-cre; Ai14* mice showing most FOS^+^ cells locate in the outlined nTS region and are not tdTomato^+^. There were strong tdTomato signals in adjoining DMV and 12N regions. Outlines in a and b indicate nTS regions at different Bregma. Scale bars, 200 µm.

**Extended Data Fig. 3.**
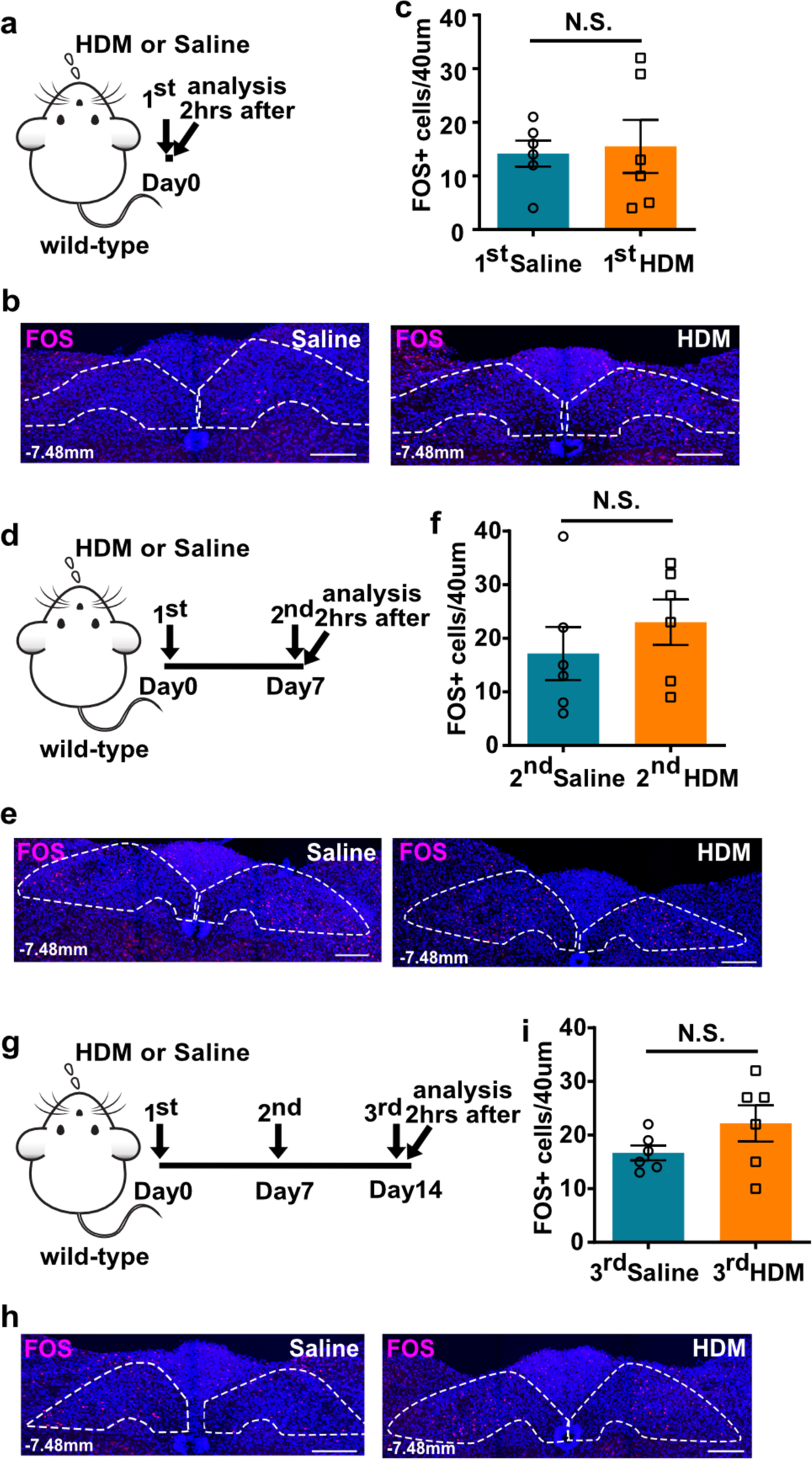
nTS neurons were activated only after repeated allergen challenge in lung. (**a**, **d**, **g**) Experiment scheme for the 1^st^, 2^nd^, or 3^rd^ HDM challenge in wild-type mice. (**b**, **c**, **e**, **f**, **h**, **i**) FOS antibody staining (b, e, h) and quantification (c, f, i) showing the lack of statistically significant increase in FOS^+^ cells after the 1^st^, 2^nd^, or 3^rd^ HDM. Outlines in b, e and h indicate nTS region at Bregma -7.48mm. Scale bars, 200 µm. Unpaired student’s *t* test was used for c, f, i. Error bars represent means ± SEM. N.S., not significant, P ≥ 0.05.

**Extended Data Fig. 4.**
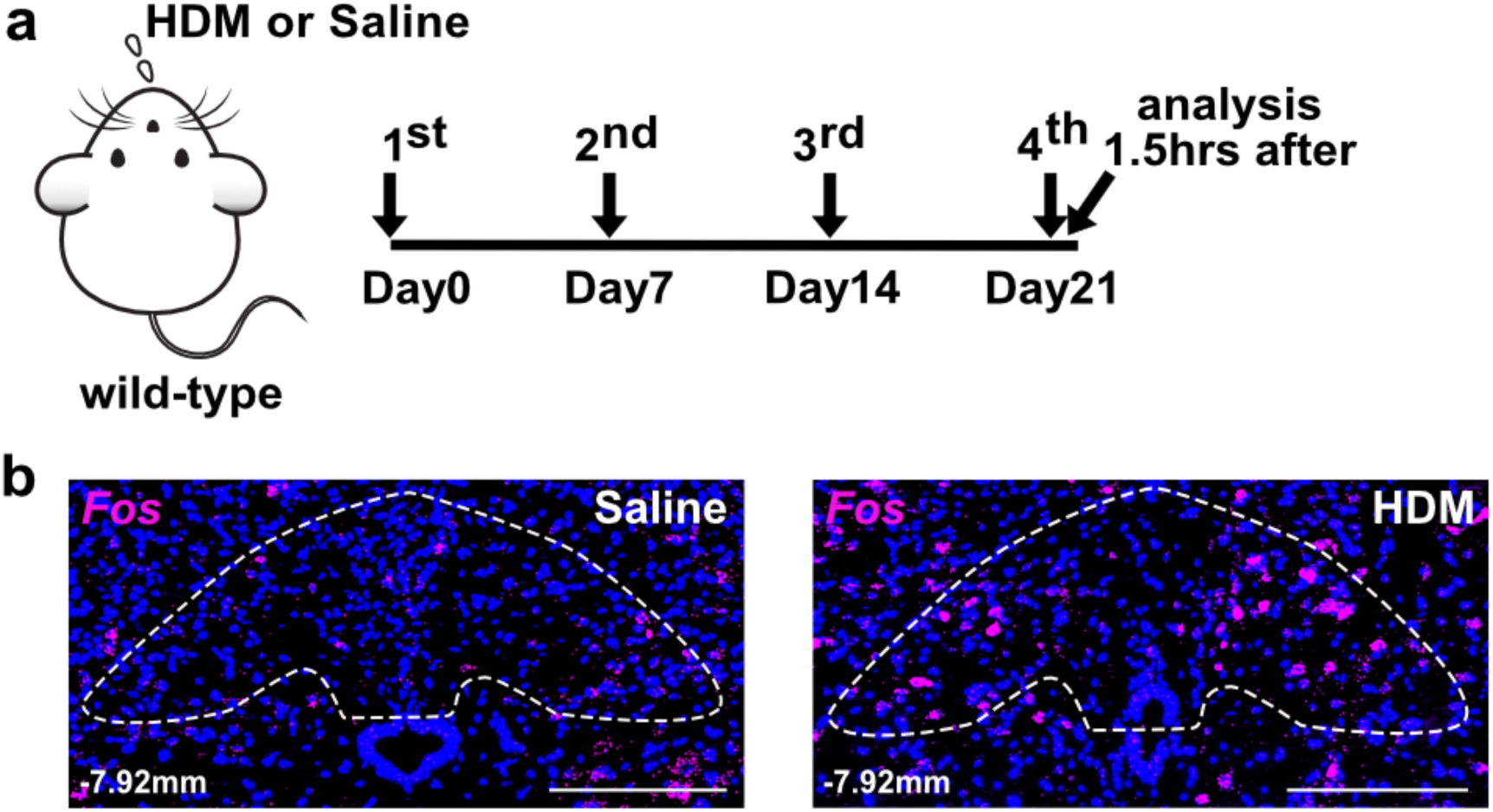
*Fos in situ* hybridization shows increased signals after allergen challenge in lung. (**a**) Experiment scheme for HDM challenge and tissue collection in wild-type mice for *Fos in situ* hybridization. (**b**) *Fos* RNA signal in the nTS is increased in the HDM group, compared to saline control. Outlines in B indicate nTS region at Bregma -7.92mm. Scale bars, 200 µm.

**Extended Data Fig. 5.**
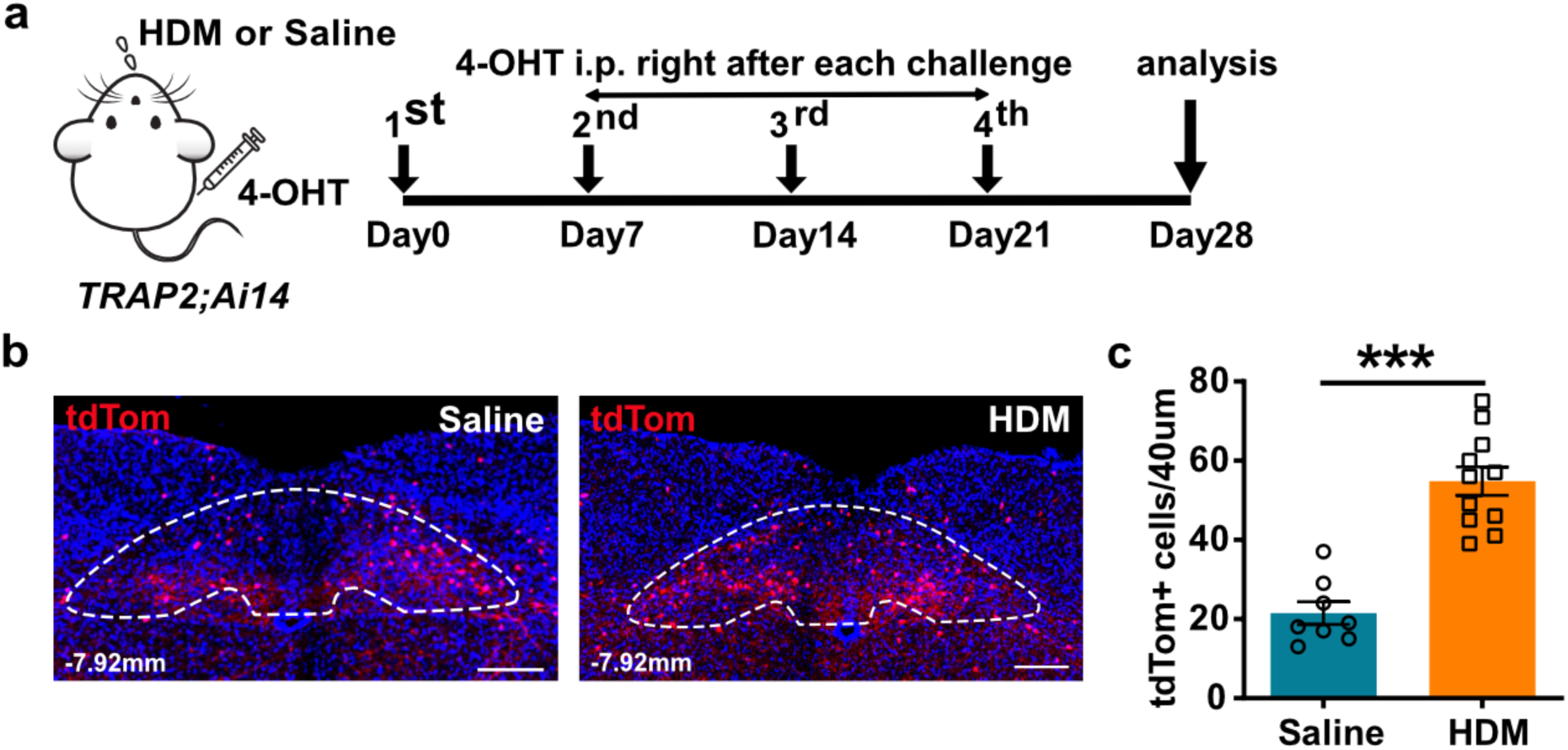
Serial sectioning and quantification of *TRAP2; Ai14* mice nTS confirmed an increase of activated tdTomato^+^ neurons in the nTS of HDM-treated mice compared to saline-treated controls. (**a**) Experiment scheme for labeling HDM-activated neurons in *TRAP2; Ai14* mice. (**b**, **c**) Representative image (b) and quantification (c) from sections of *TRAP2; Ai14* mice showing increased tdTomato signals in the nTS after HDM at Bregma -7.92 mm. Outlines in b indicate nTS region. Scale bars, 200 µm. Unpaired student’s *t* test was used for c. Error bars represent means ± SEM, ***p<0.001.

**Extended Data Fig. 6.**
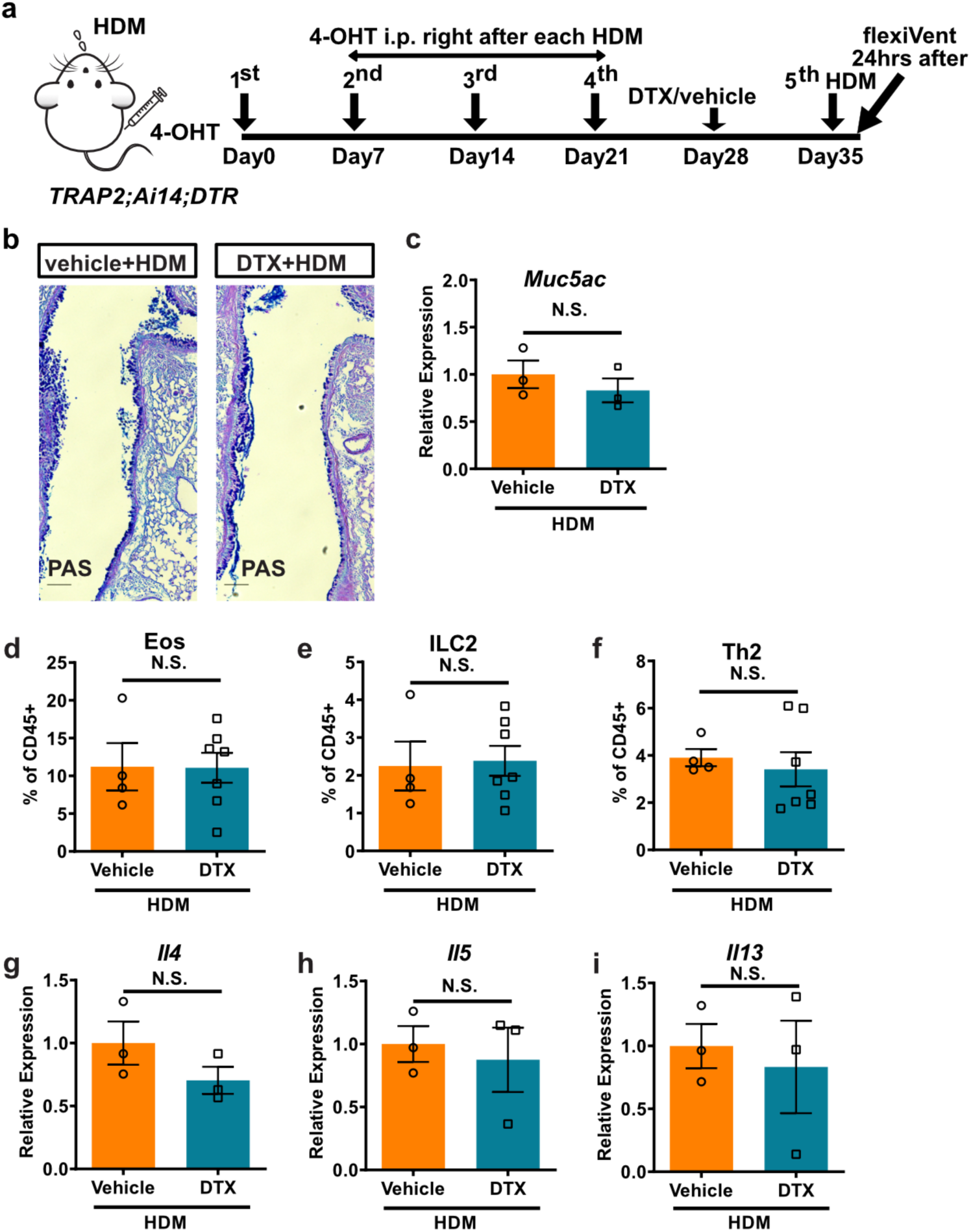
Genetic ablation of HDM-activated neurons in the nTS did not affect goblet cell metaplasia or immune cell infiltration. (**a**) Experimental scheme for DTX ablation of HDM-activated nTS neurons in *TRAP2; Ai14; DTR* mice. (**b**, **c**) PAS staining (b) and qPCR of *Muc5ac* (c) showing no change in HDM-induced goblet cell metaplasia response between nTS vehicle-injected and DTX-injected *TRAP2; Ai14; DTR* mice. (**d**-**i**) Flow cytometry analyses of eosinophils (Eos, d), innate lymphoid cells (ILC2s, e) and T-helper type 2 cells (Th2, f), and qPCR of *Il4* (g), *Il5* (h) and *Il13* (i) from whole lungs of vehicle-injected or DTX-injected mice showing no statistically significant differences between groups. Scale bars, 100 µm in b. Unpaired student’s *t* test was used for c-i. Error bars represent means ± SEM. N.S., not significant.

**Extended Data Fig. 7.**
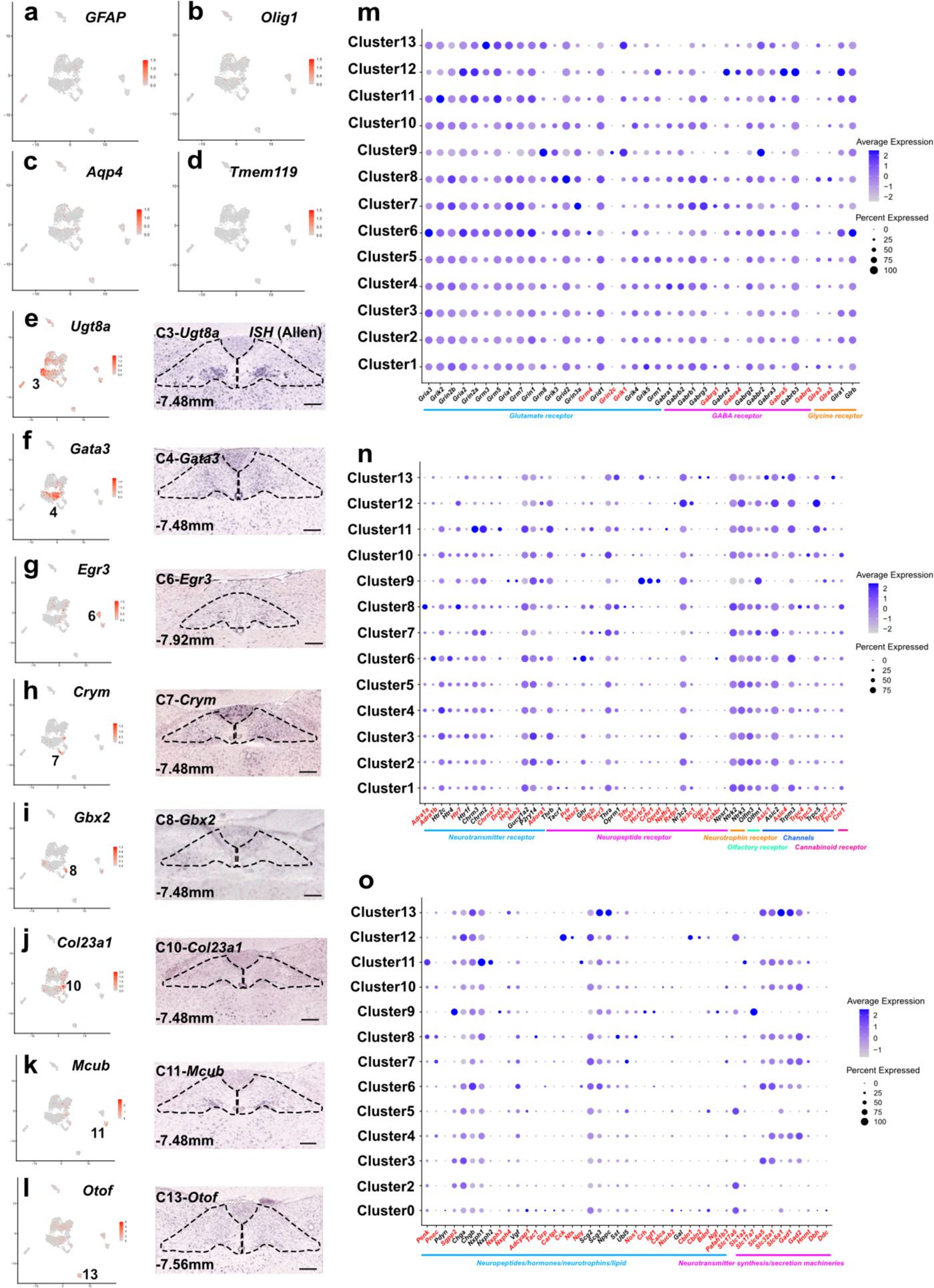
snRNA-seq provided a transcriptomic atlas of nTS. (**a**-**d**) UMAP plots showing little to no expression of glia marker genes-*GFAP* (a), *Olig1* (b), *Aqp4* (c) or *Tmem119* (d) in the neuron clusters. (**e**-**l**) UMAP plots (left) and RNA *in situ* hybridization images from the Allen Brain Atlas (right) showing expression of marker genes in the nTS inhibitory clusters as labeled. Genes plotted here were indicated in red in Fig. 2g. Scale bars, 100 µm for e-l. (**m**-**o**) Dot plot showing nTS neuron expression of factors involved in neuronal signal reception and transmission. Gene names in red are ones with notable varying expression across the 13 nTS clusters.

**Extended Data Fig. 8.**
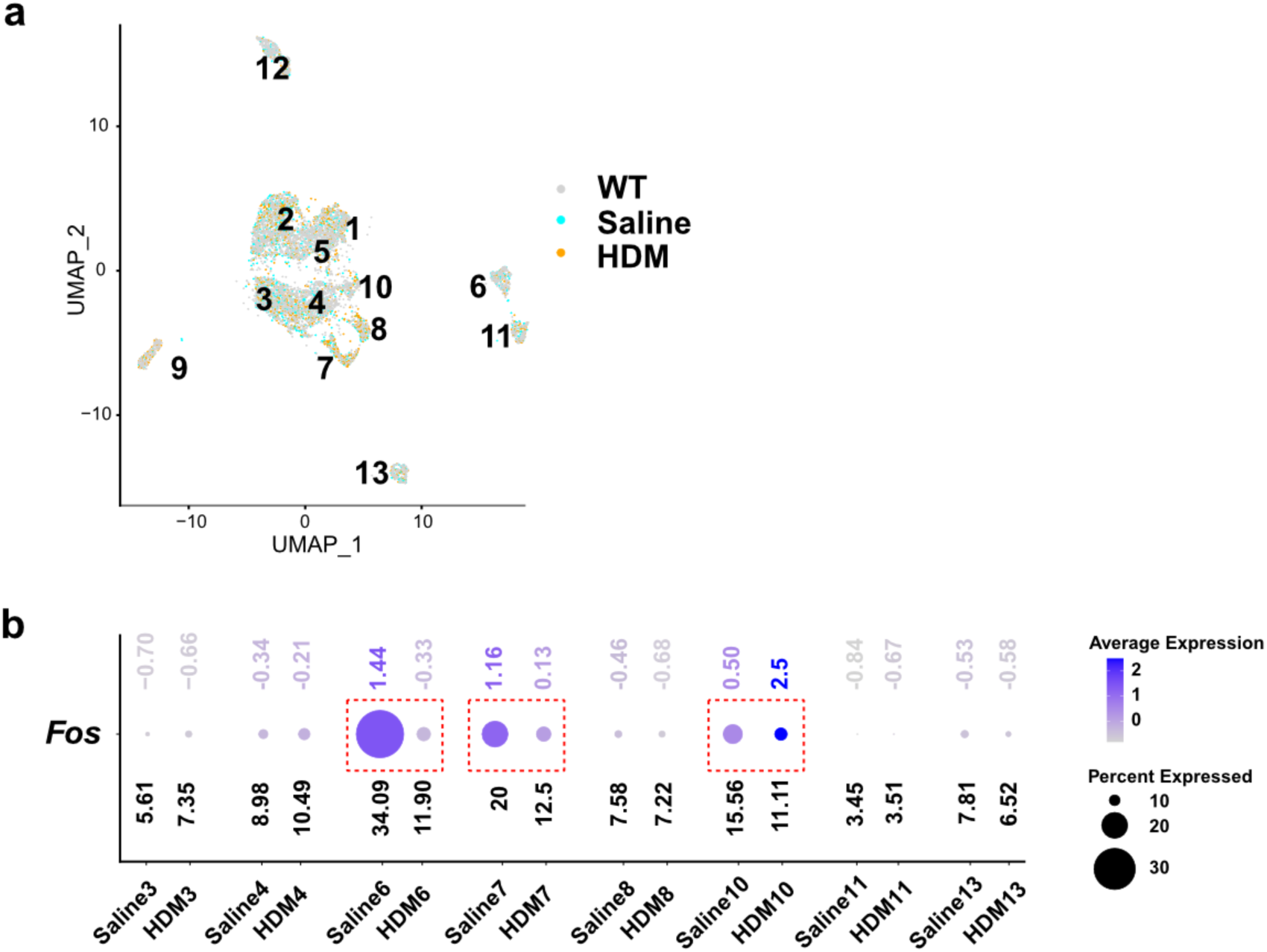
*Dbh^+^* neurons in the nTS were activated upon allergen challenge in lung. (**a**) UMAP plot pseudo-colored to show representation of neurons in all 13 nTS clusters from wild-type, saline-treated or HDM-treated mice. (**b**) Dot plot of *Fos* expression in individual inhibitory nTS clusters after either saline or HDM challenge to the lung. Numbers on upper row denote Log_2_ average expression levels, and numbers on bottom row denote percentages of expressing cells. Clusters 6, 7 and 10 populations (boxed) show decreased proportions after HDM challenge compared to saline control.

**Extended Data Fig. 9.**
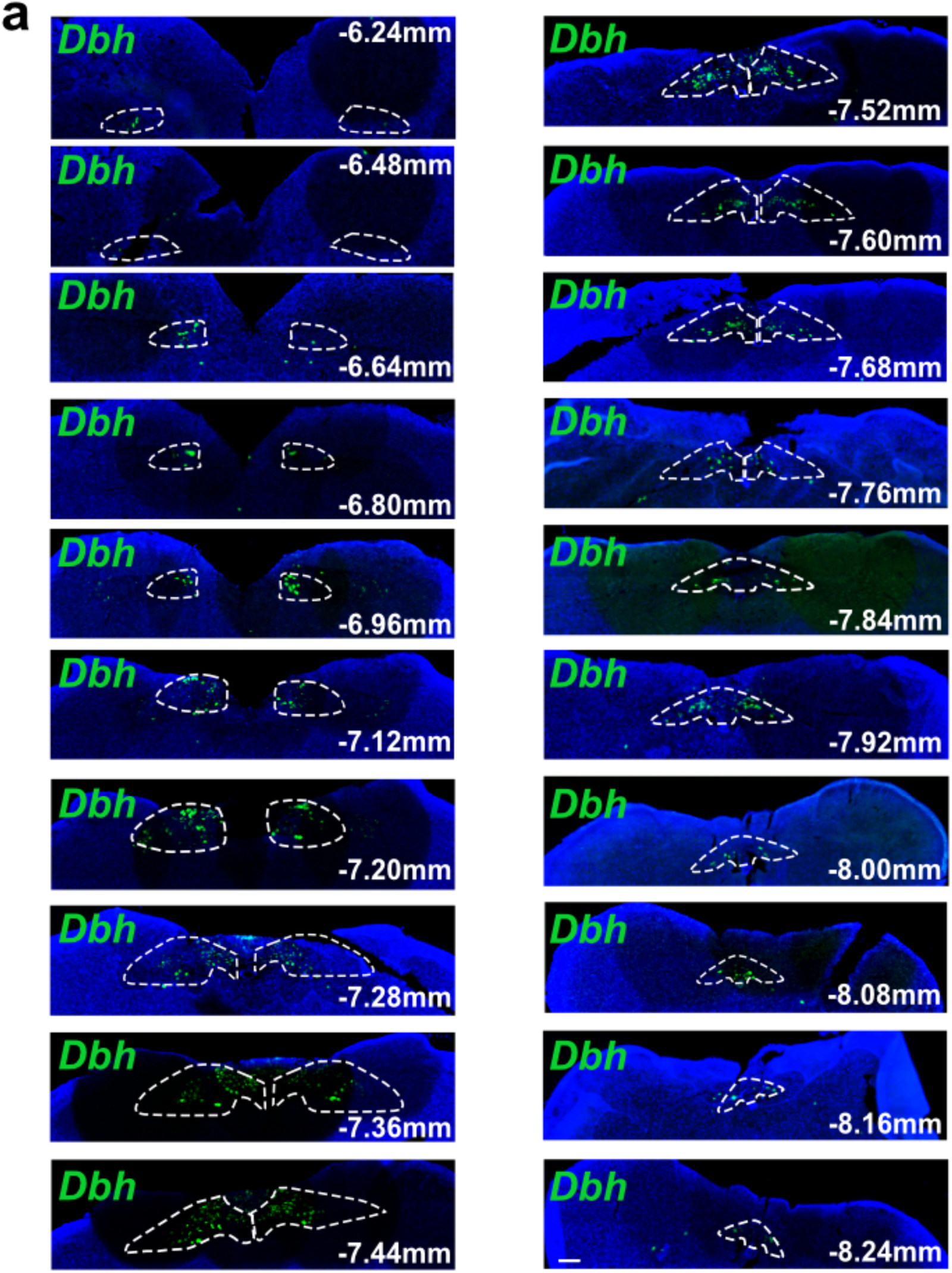
Expression pattern of *Dbh* in the whole nTS. (**a**) *Dbh* RNAscope *in situ* hybridization on serial sections of naïve brainstem that covers most of the whole nTS regions from Bregma -6.96 mm to Bregma -8.08 mm. Scale bar, 200 µm.

**Extended Data Fig. 10.**
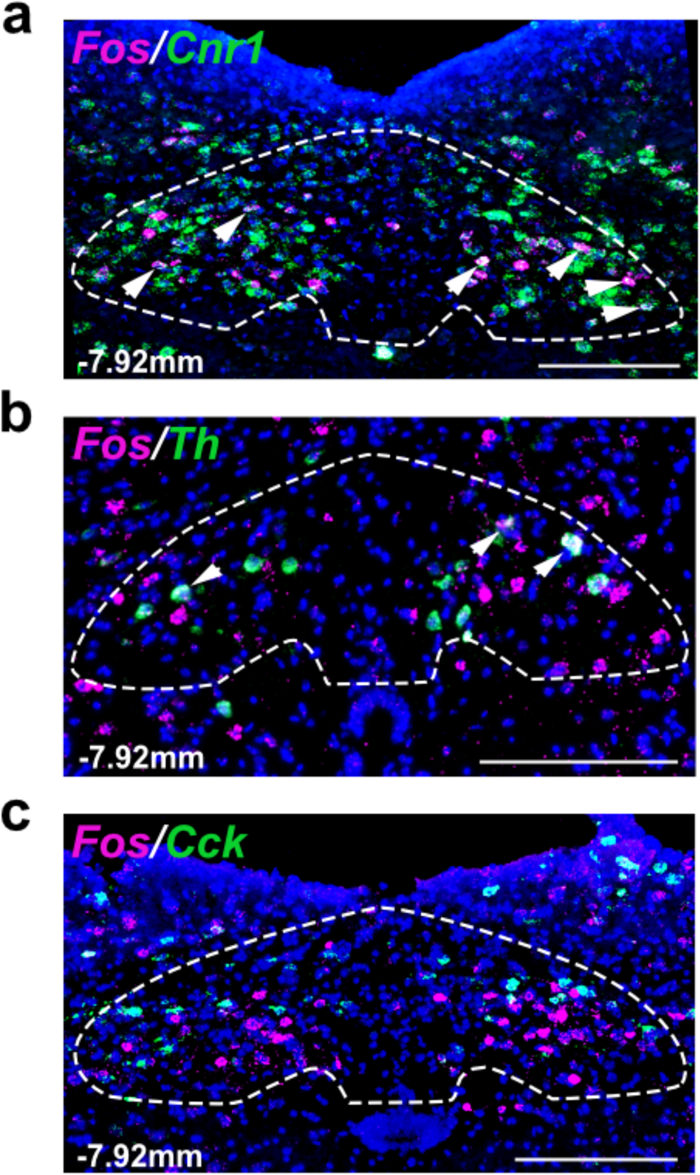
Double *in situ* hybridization between *Fos* and *Cnr1*, or *Th*, or *Cck* after HDM challenge. (**a**-**c**) Double RNAscope *in situ* hybridization showing some overlap between *Fos* and *Cnr1* (a) or *Th* (b), and little to no overlap between *Fos* and *Cck* (c) in the nTS at 1.5 hours after HDM challenge to lung. Outlines in a-c indicate nTS region at Bregma -7.92 mm. Scale bars, 200 µm.

**Extended Data Fig. 11.**
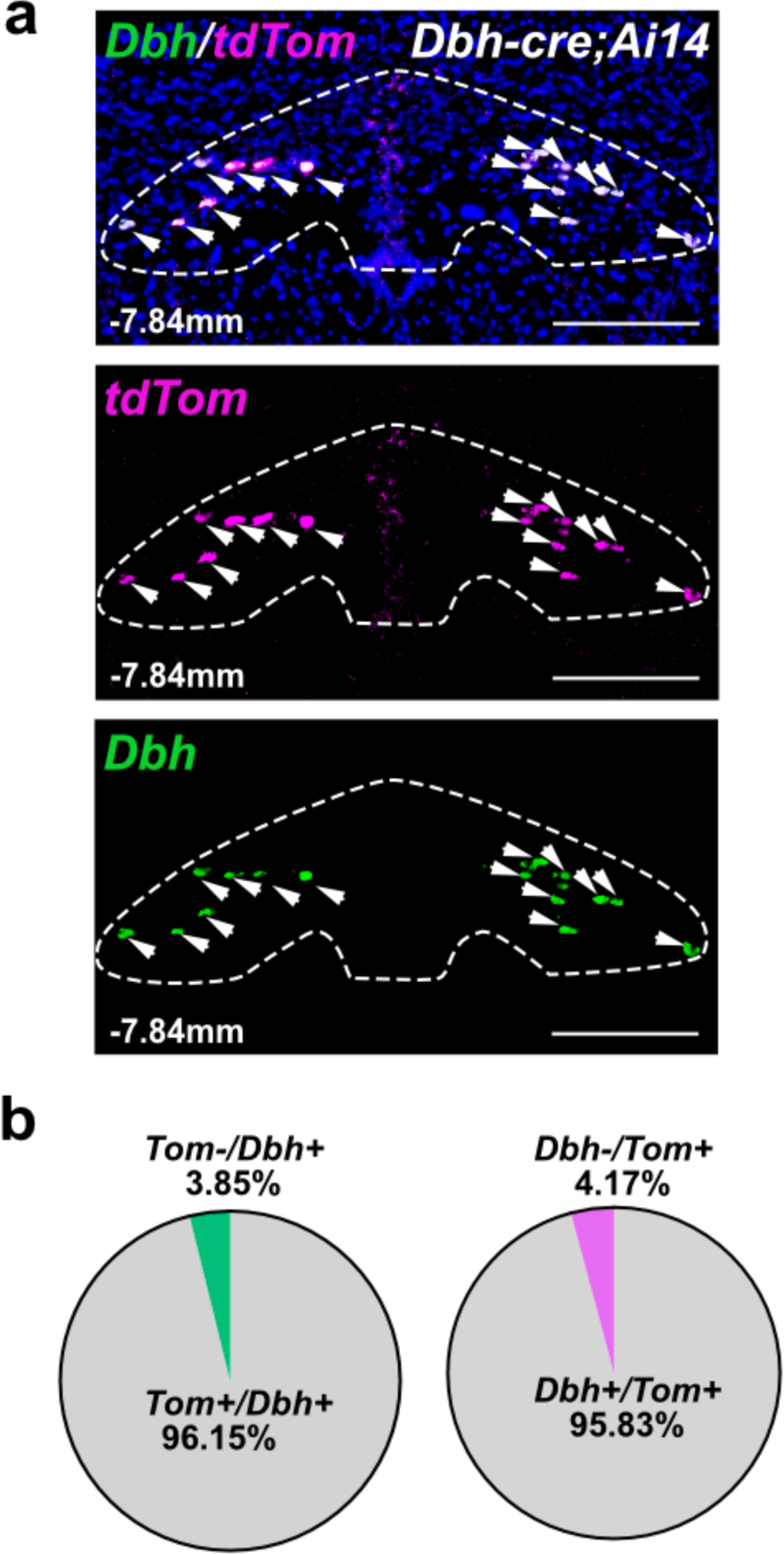
*Dbh^+^*neurons are genuinely captured by *Dbh-cre; tdTom* labeling. (**a**, **b**) Double *in situ* hybridization representative images (a) and quantification (b) showing high overlap between *Dbh* and *tdTomato* in the nTS of *Dbh-cre; tdTomato* mice. Outlines in a indicate nTS region at Bregma -7.84 mm. Arrowheads in a indicate cells with overlapped signals. Scale bars, 200 µm.

**Extended Data Fig. 12.**
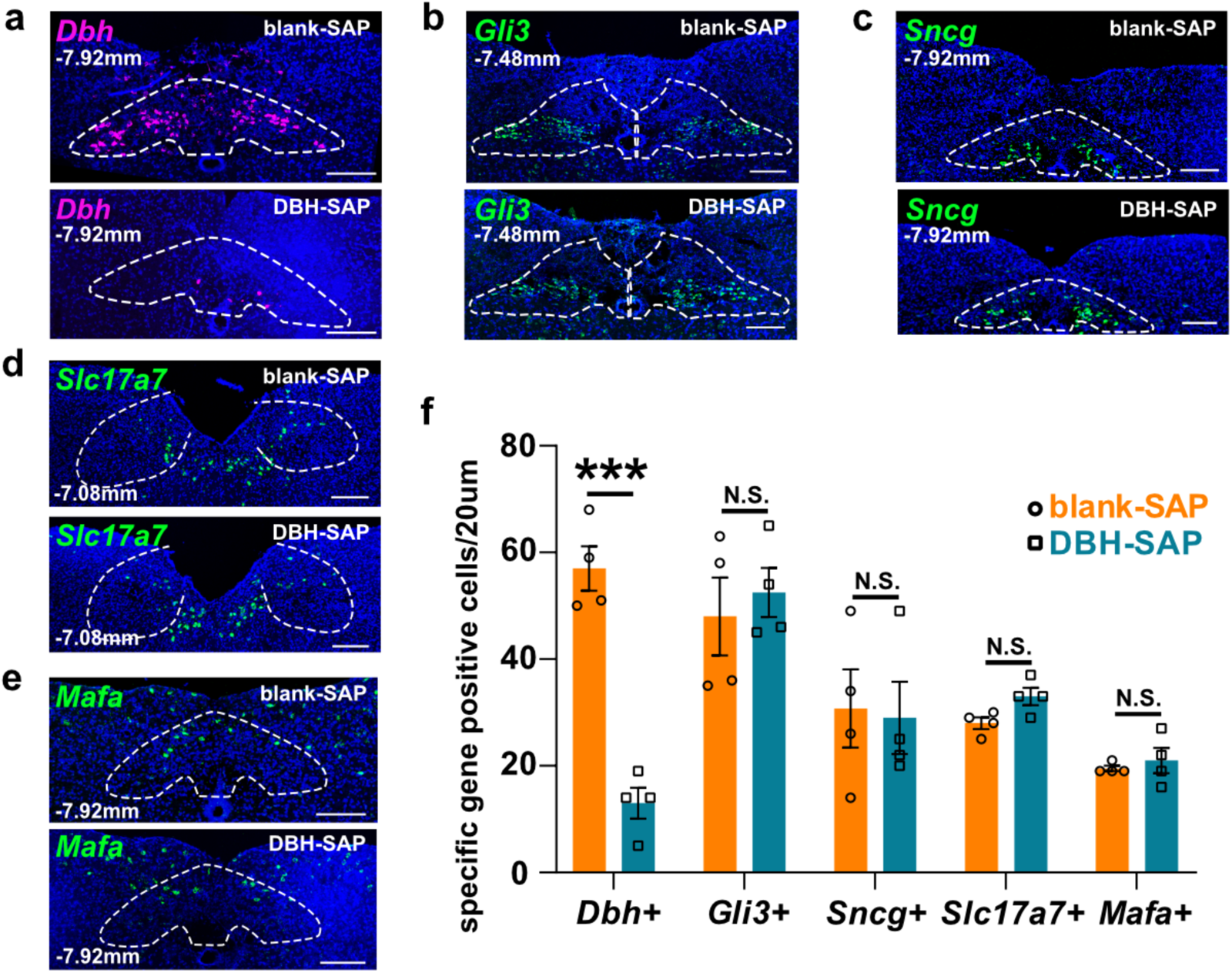
No change in the expression of other excitatory nTS neuronal subsets after anti-DBH-SAP injection into nTS. (**a**-**f**) Representative *in situ* hybridization of nTS excitatory cluster marker genes *Dbh*, *Gli3*, *Sncg*, *Slc17a7*, *Mafa* (a-e) and quantification (f, each datapoint represents the average number from one mouse) showing positive cells in nTS region following DBH-SAP bilateral injection into wild type nTS, compared to blank-SAP treatment. Outlines in a-e indicate nTS regions at different Bregma. Scale bars, 200 µm in a-e. Unpaired student’s *t* test was used for f. Error bars represent means ± SEM. N.S., not significant, P ≥ 0.05; ***P < 0.001.

**Extended Data Fig. 13.**
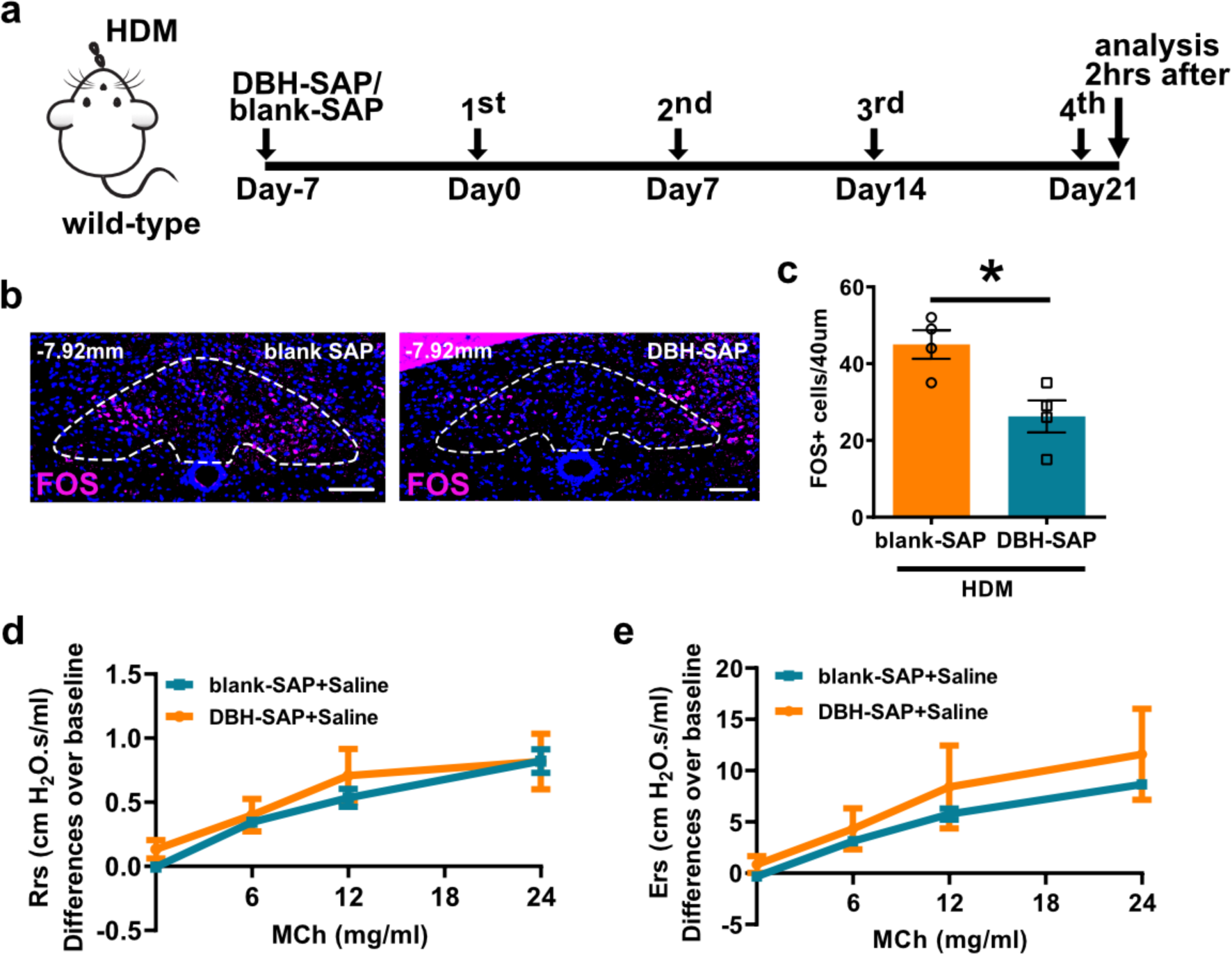
Chemical depletion of *Dbh*^+^ neurons in the nTS reduced FOS^+^ cells after HDM challenge in lung while had no effect on airway hyperreactivity on naïve mice. (**a**) Experiment scheme for detection of FOS^+^ cells after DBH-SAP treatment in wild-type mice. (**b**, **c**) Representative staining (b) and quantification (c) showing reduced FOS^+^ cells after ablation of *Dbh^+^* neurons in the nTS following DBH-SAP treatment scheme. (**d**, **e**) FlexiVent data showing that DBH-SAP bilateral injection into nTS of naïve mice had no effect on airway constriction in response to increasing doses of methacholine. Outlines in b indicate nTS region at Bregma -7.92 mm. Scale bars, 200 µm in b. Unpaired student’s *t* test was used for c, two-way ANOVA was used for d, e, statistical analysis was performed separately at each MCh concentration. Error bars represent means ± SEM, *P < 0.05; N.S., not significant, P ≥ 0.05.

**Extended Data Fig. 14.**
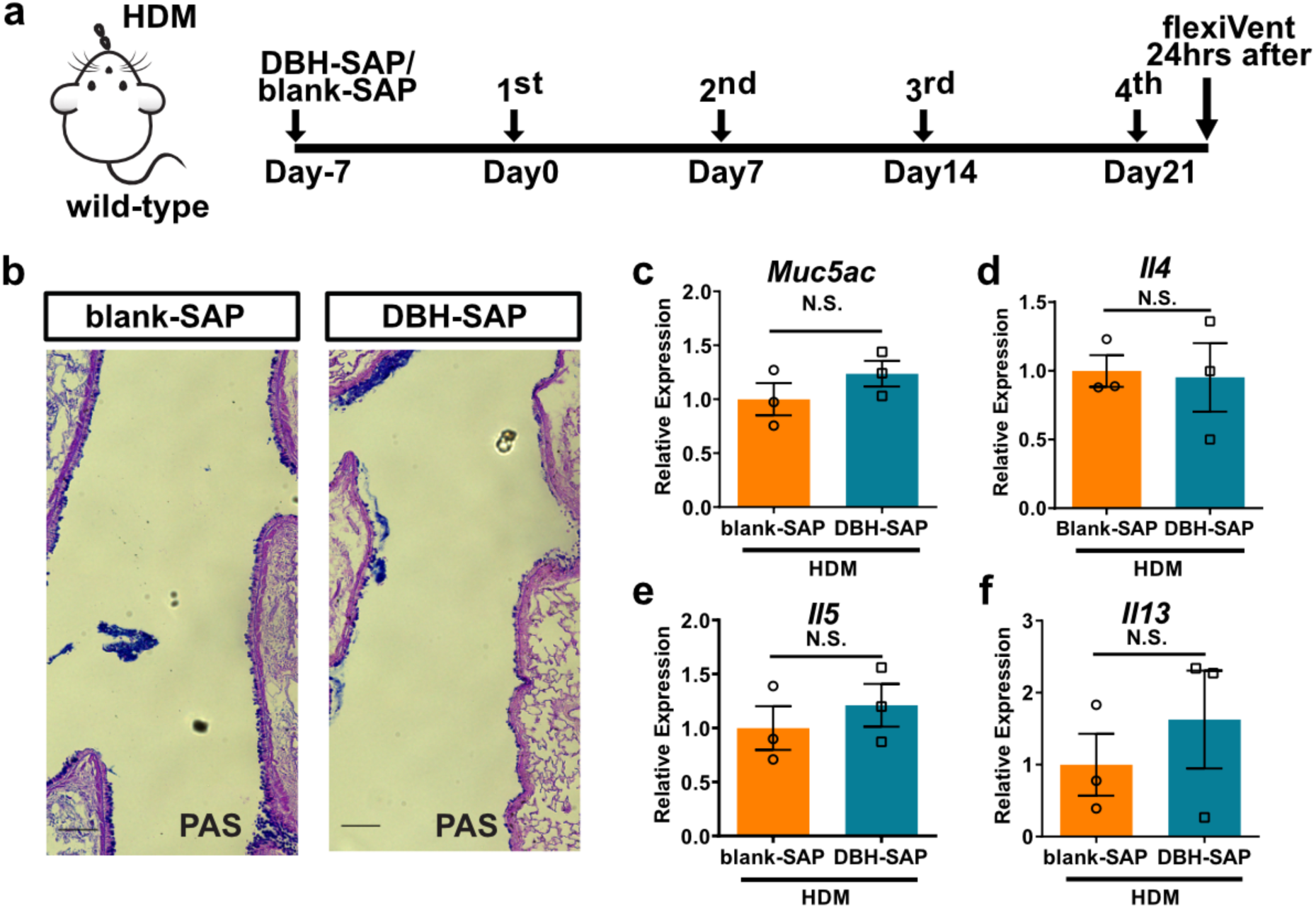
Chemical depletion of *Dbh*^+^ neurons did not affect allergen-induced goblet cell metaplasia or cytokine genes expressions. (**a**) Experiment scheme for flexiVent measurement after DBH-SAP treatment in wild-type mice. (**b**) PAS staining showing no change in HDM-induced goblet cell metaplasia response between blank-SAP-injected and DBH-SAP-injected wild-type mice. (**c**-**f**) Whole lung qPCR of goblet cell marker *Muc5ac* (c) and key type 2 immunity cytokine genes *Il4* (d), *Il5* (e) and *Il13* (f) showing no statistic difference between blank-SAP-injected and anti-DBH-SAP-injected wild-type mice. Scale bars, 100 µm in b. Unpaired student’s *t* test was used for c-f. Error bars represent means ± SEM. N.S., not significant, P ≥ 0.05.

**Extended Data Fig. 15.**
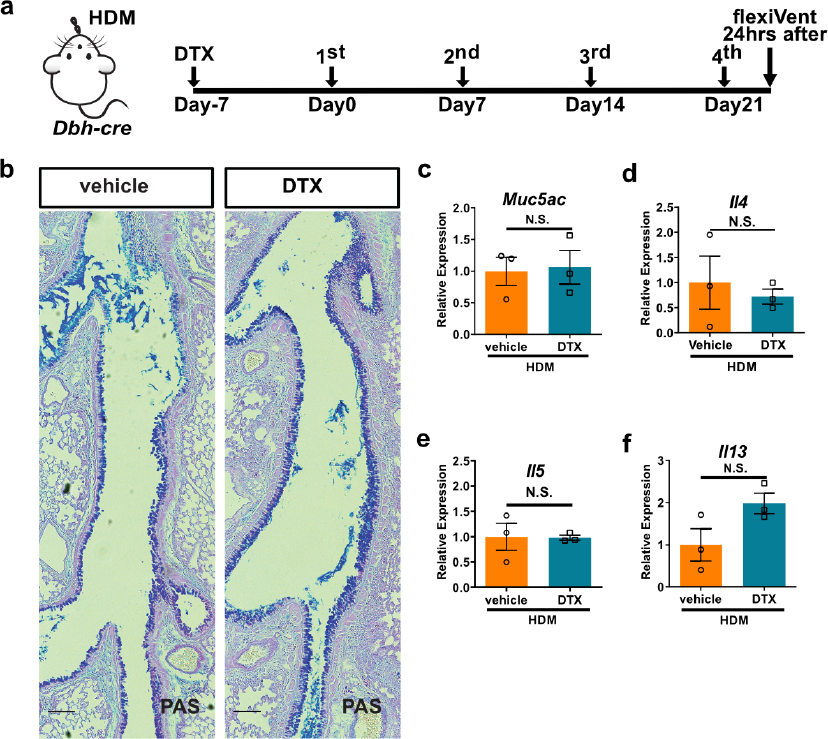
Genetic depletion of *Dbh*^+^ neurons did not affect allergen-induced goblet cell metaplasia or cytokine gene expressions. (**a**) Experiment scheme for DTX injection on *Dbh-cre; DTR* mice. (**b**) PAS staining showed no change in HDM-induced goblet cell metaplasia response between vehicle-and DTX-injected *Dbh-cre; DTR* mice. (**c**-**f**) qPCR of goblet cell marker *Muc5ac* (c), key type 2 immunity cytokines *Il4* (d), *Il5* (e) and *Il13* (f) in vehicle-or DTX-injected *Dbh-cre; DTR* mice. Scale bars, 100 µm in b. Unpaired student’s *t* test was used for c-f. Error bars represent means ± SEM. N.S., not significant.

**Extended Data Fig. 16.**
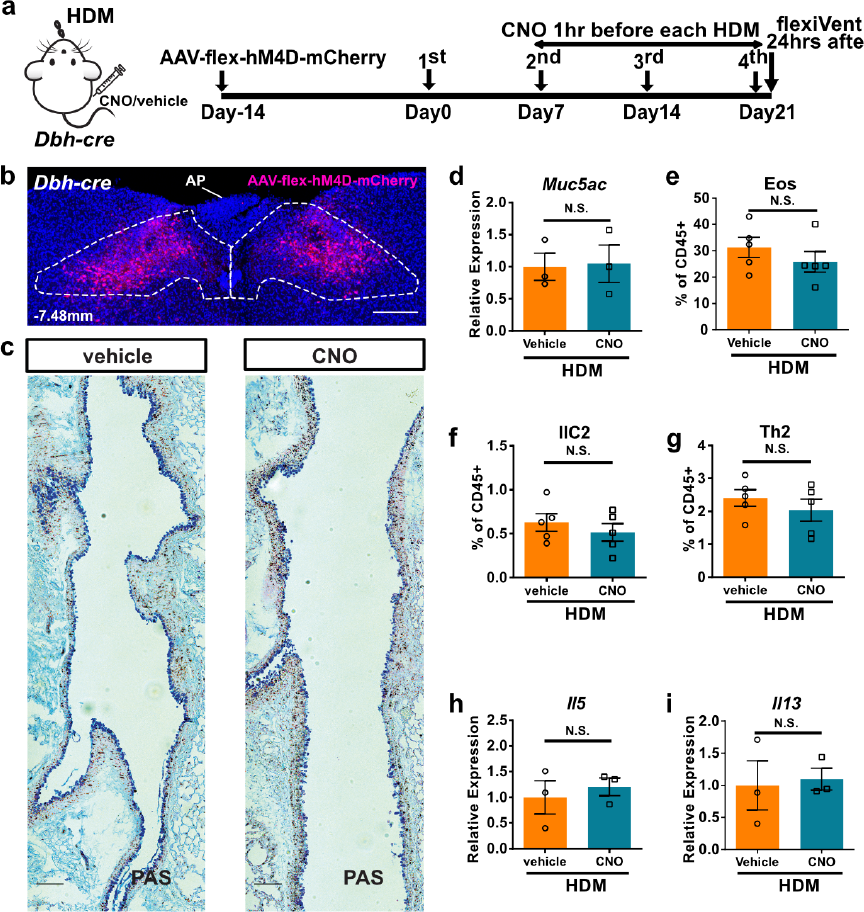
Genetic inactivation of *Dbh*^+^ neurons in the nTS did not affect allergen-induced goblet cell metaplasia or immune cell infiltration. (**a**) Experiment scheme for *Dbh^+^* neuron inhibition following AAV-flex-hM4D-mCherry injection in *Dbh-cre* mice. (**b**) Representative mCherry signals (with Dsred antibody staining, at Bregma -7.48 mm) following bilateral AAV-hM4D-mCherry injection in the nTS (but not AP). (**c**, **d**) PAS staining (c) and qPCR of *Muc5ac* (d) showed no change in HDM-induced goblet cell metaplasia response between vehicle-and CNO-injected lungs after AAV-flex-hM4D-mCherry injection into bilateral nTS. (**e**-**i**) Flow cytometry analyses of immune cells (e-g), and qPCR of *Il5* (h) and *Il13* (i) from whole lungs showing no change in HDM-induced immune cell infiltration between vehicle- and CNO-injected lungs after AAV-flex-hM4D-mCherry injection into bilateral nTS. Outlines in b indicate dorsomedial part of nTS region at Bregma -7.48 mm. Scale bars, 200 µm in b, 100 µm in c. Unpaired student’s *t* test was used for d-i. Error bars represent means ± SEM. N.S., not significant.

**Extended Data Fig. 17.**
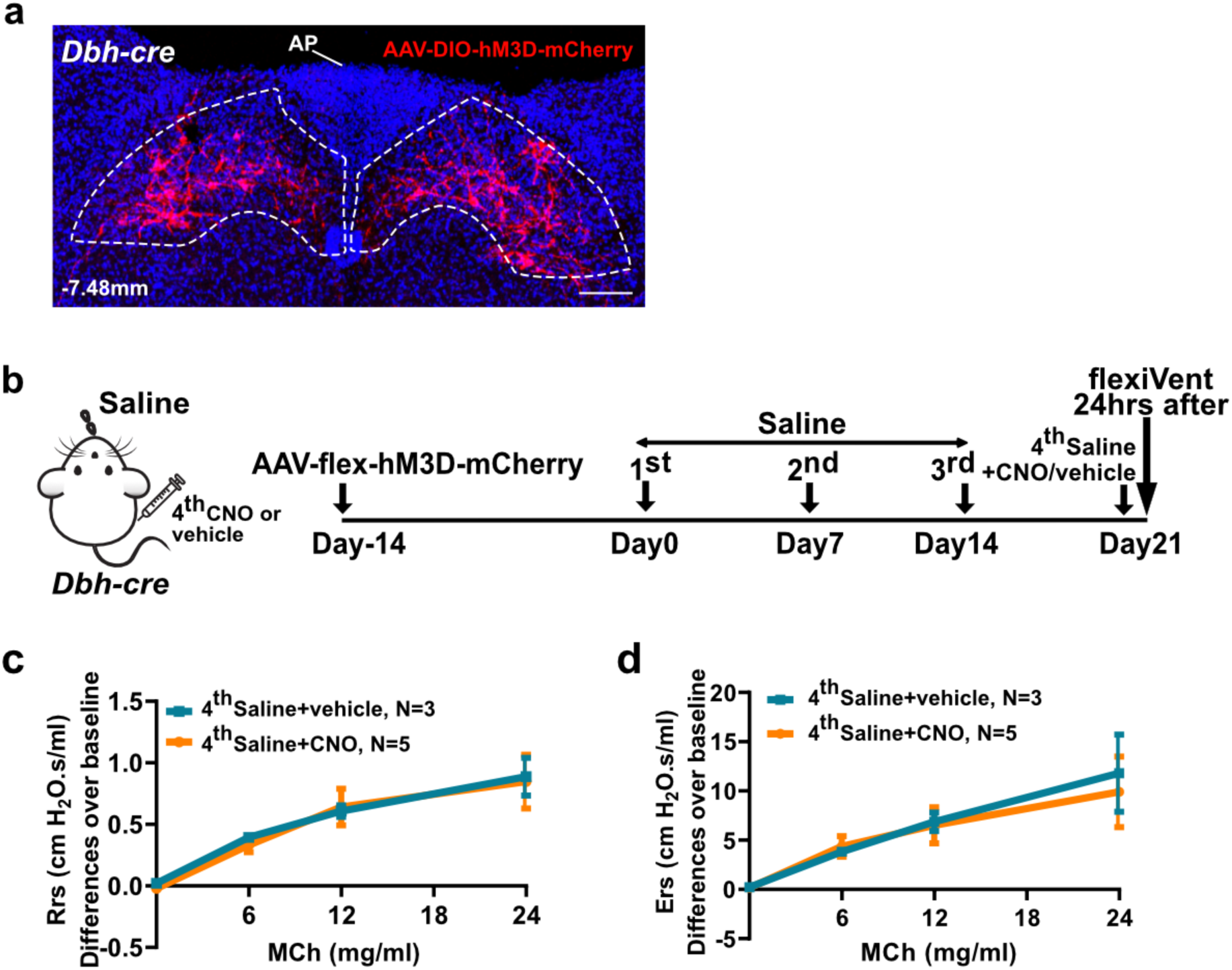
Genetic activation of *Dbh*^+^ neurons in the nTS had no effect on airway hyperreactivity on naïve mice. (**a**) Representative mCherry signals (with Dsred antibody staining) following bilateral AAV-DIO-hM3D-mCherry injection in the nTS (but not AP). (**b**) Experiment scheme for CNO activation of nTS *Dbh^+^* neurons on naïve mice. Both groups received the 1^st^-3^rd^ saline challenge. (**c**, **d**) FlexiVent data showing that CNO activation (4^th^ dose) of *Dbh^+^*nTS neurons had no effect on airway constriction in response to increasing doses of methacholine. Outlines in a indicate nTS region at Bregma -7.48 mm. Scale bars, 200 µm in a. Two-way ANOVA was used for c, d, statistical analysis was performed separately at each MCh concentration. Error bars represent means ± SEM. N.S., not significant, P ≥ 0.05.

**Extended Data Fig. 18.**
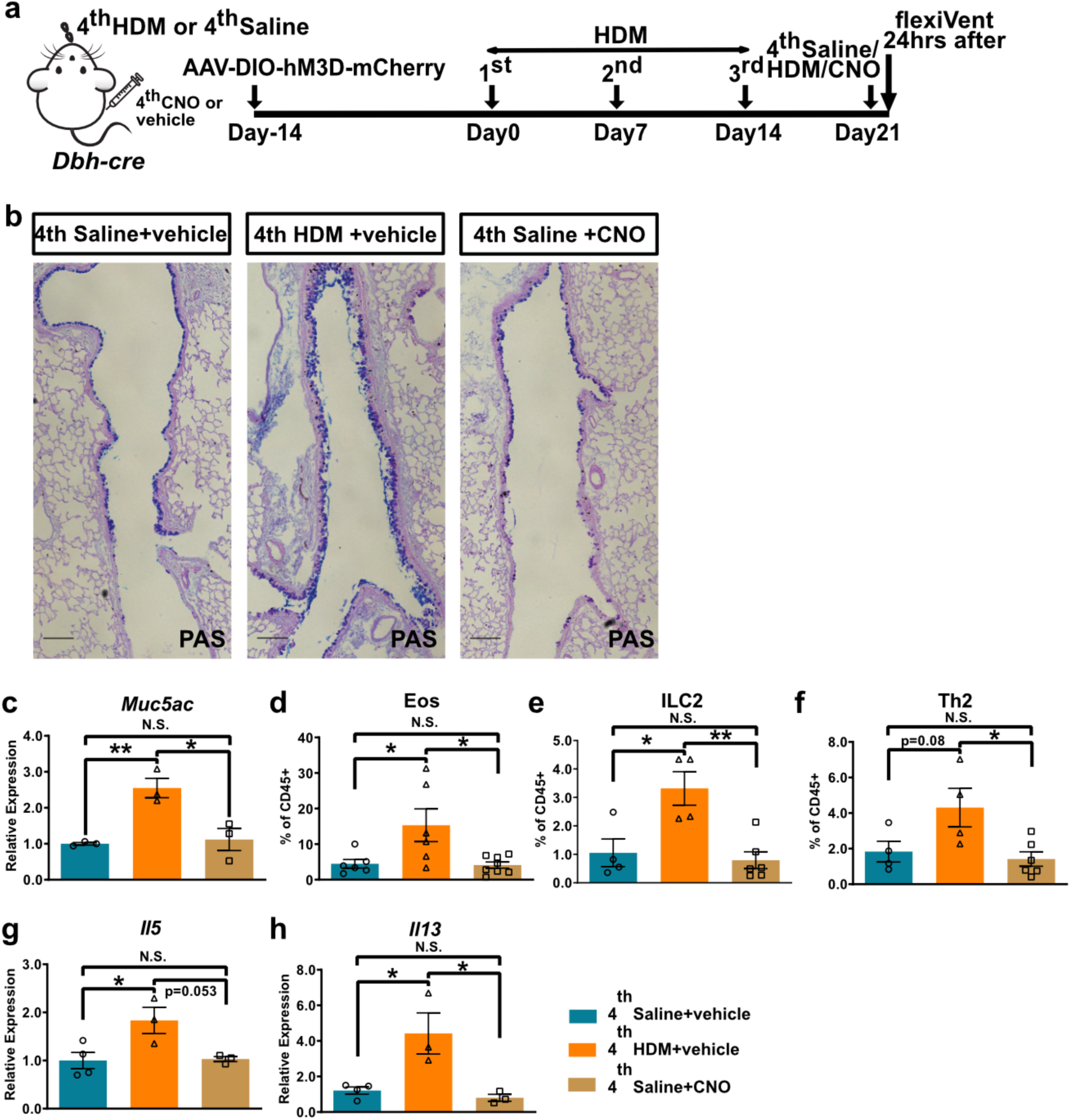
Genetic activation of *Dbh*^+^ neurons in the nTS did not affect allergen-induced goblet cell metaplasia or immune cell infiltration. (**a**) Experiment scheme for CNO activation of nTS *Dbh^+^* neurons on sensitized mice. Both groups received the 1^st^-3^rd^ HDM challenge. (**b**, **c**) PAS staining (b) and qPCR of *Muc5ac* (c) from whole lungs showing no change in HDM-induced goblet cell metaplasia response following 4^th^ *Dbh^+^* neuronal activation in the nTS compared to 4^th^ saline controls. (**d**-**h**) Flow cytometry analyses of immune cells (d-f), and qPCR of *Il5* (g) and *Il13* (h) from whole lungs showing no change in HDM-induced immune cell infiltration following 4^th^ *Dbh^+^* neuronal activation in the nTS compared to 4^th^ saline controls. Scale bars, 100 µm in b. One-way ANOVA was used for c-h. Error bars represent means ± SEM. N.S., not significant, P ≥ 0.05; *P < 0.05; **P < 0.01.

**Extended Data Fig. 19.**
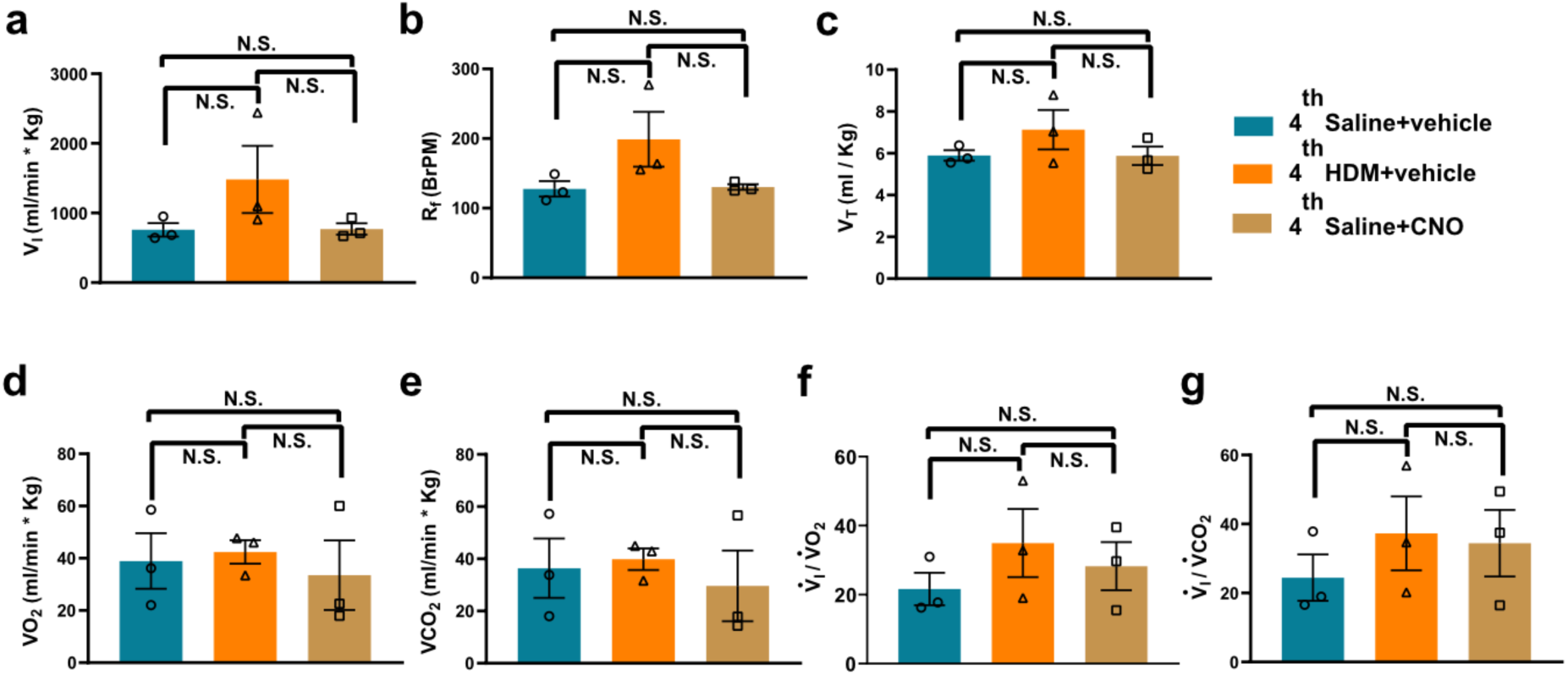
Genetic activation of *Dbh*^+^ neurons in the nTS did not change respiratory parameters in response to normal levels of O_2_ and CO_2_, as measured by plethysmograph. (**a**-**g**) Measurements of ventilation showing no change in respiratory parameters including minute ventilation (V_I_, a) , respiratory frequency (R_f_, b) and tidal volume (V_t_, c), as well as measurements for metabolic rate that includes the V_O2_ (d) and V_CO2_ (e), and the “real” ventilation considering the metabolic rate (Vi/VO2, f and Vi/VCO_2_, g) across groups. VI, inspired ventilation, product of frequency and tidal volume which was normalized to body mass ventilation (ml/min·kg). VO_2_, oxygen consumption, V_CO2_, carbon dioxide. The ratio of Vi/V_O_2__ and Vi/V_CO_2__ provide a precise assessment of ventilation by avoiding confounding factors produced by changes in metabolic rate. One-way ANOVA was used for a-g. Error bars represent means ± SEM. N.S., not significant, P ≥ 0.05.

**Extended Data Fig. 20.**
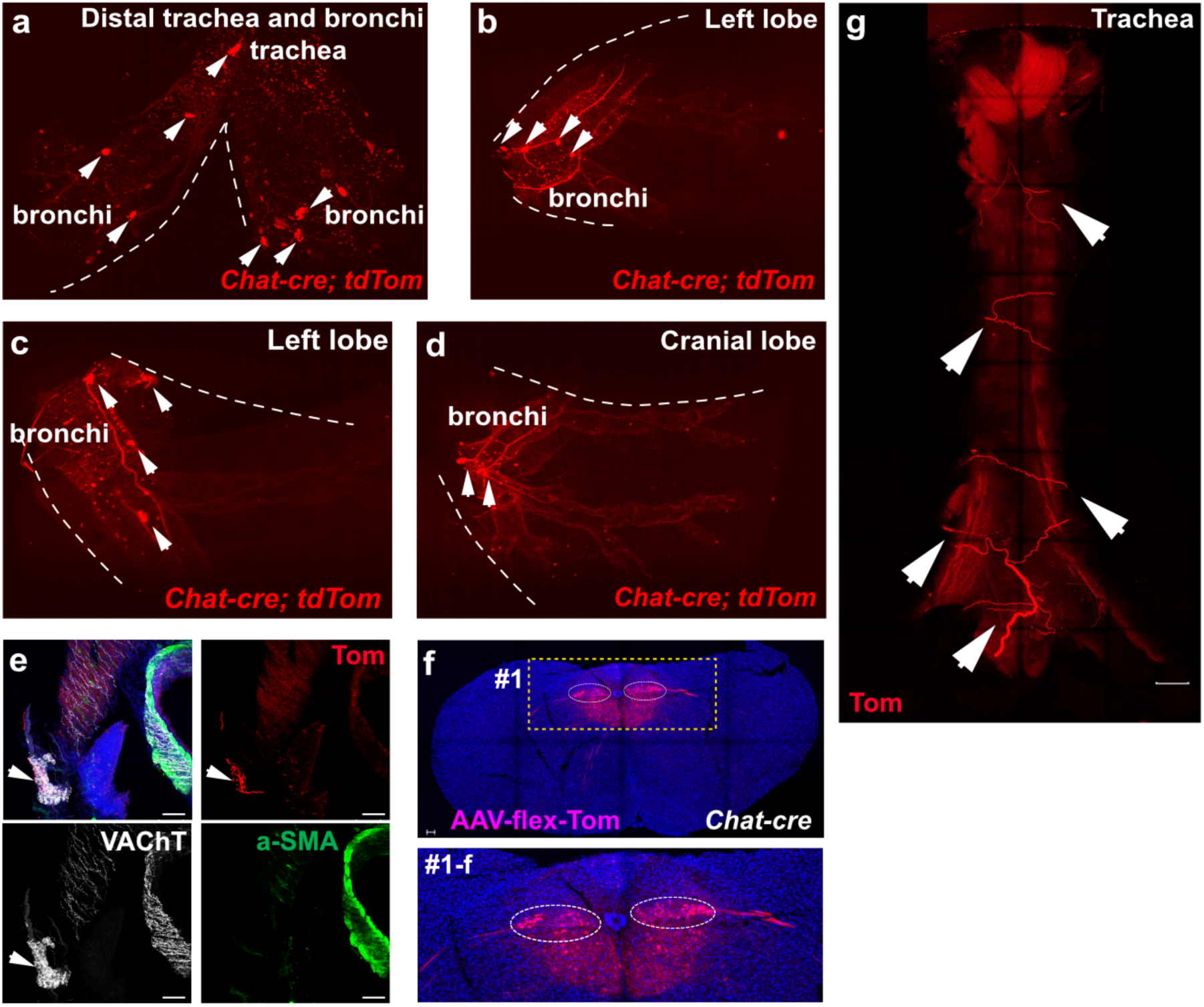
Parasympathetic neurons in the NA, but not the DMV or 12N, project to the airway-innervating postganglionic neurons. (**a**-**d**) tdTomato^+^ neurons/ganglia (arrowheads) reside in the trachea, main bronchi (a) and main airway entrance (hilum) of lung lobes (b-d) of *Chat-cre; tdTomato* mice. (**e**) tdTomato^+^ nerves from NA innervate postganglionic parasympathetic ganglia (labeled with VAChT antibody, arrowhead) in the extrapulmonary airway, rather than directly on smooth muscle cells (labeled with a-SMA antibody). (**f**) Stereotaxic injection of AAV-flex-tdTomato into bilateral DMV (also labeled neurons in ventral adjoining 12N) of *Chat-cre* mice. Boxed area in #1 was magnified below. Ovals in #1 outline DMV region. (**g**) tdTomato^+^ nerves from DMV were found passing through the space between trachea and esophagus (arrowheads). Scale bars, 100 µm in e, f, 400 µm in g.

**Extended Data Fig. 21.**
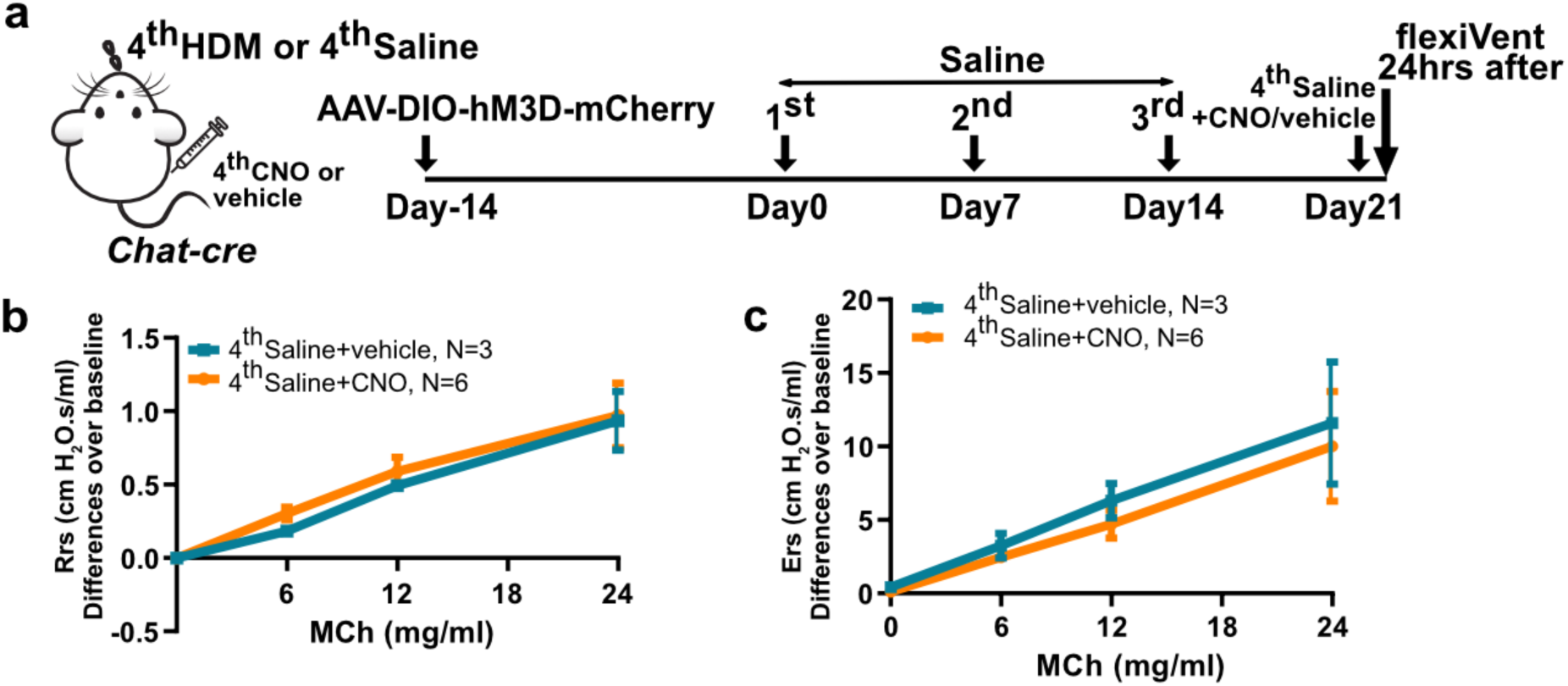
Genetic activation of *Chat*^+^ neurons in the NA had no effect on airway hyperreactivity on naïve mice. (**a**) Experiment scheme for CNO activation of NA *Chat^+^* neurons on naïve mice. Both groups received the 1^st^-3^rd^ saline challenge. (**b**, **c**) FlexiVent data showing that CNO activation (4^th^ dose) of *Chat^+^*NA neurons had no effect on airway constriction in response to increasing doses of methacholine. Two-way ANOVA was used for b, c, statistical analysis was performed separately at each MCh concentration. Error bars represent means ± SEM. N.S., not significant, P ≥ 0.05.

**Extended Data Fig. 22.**
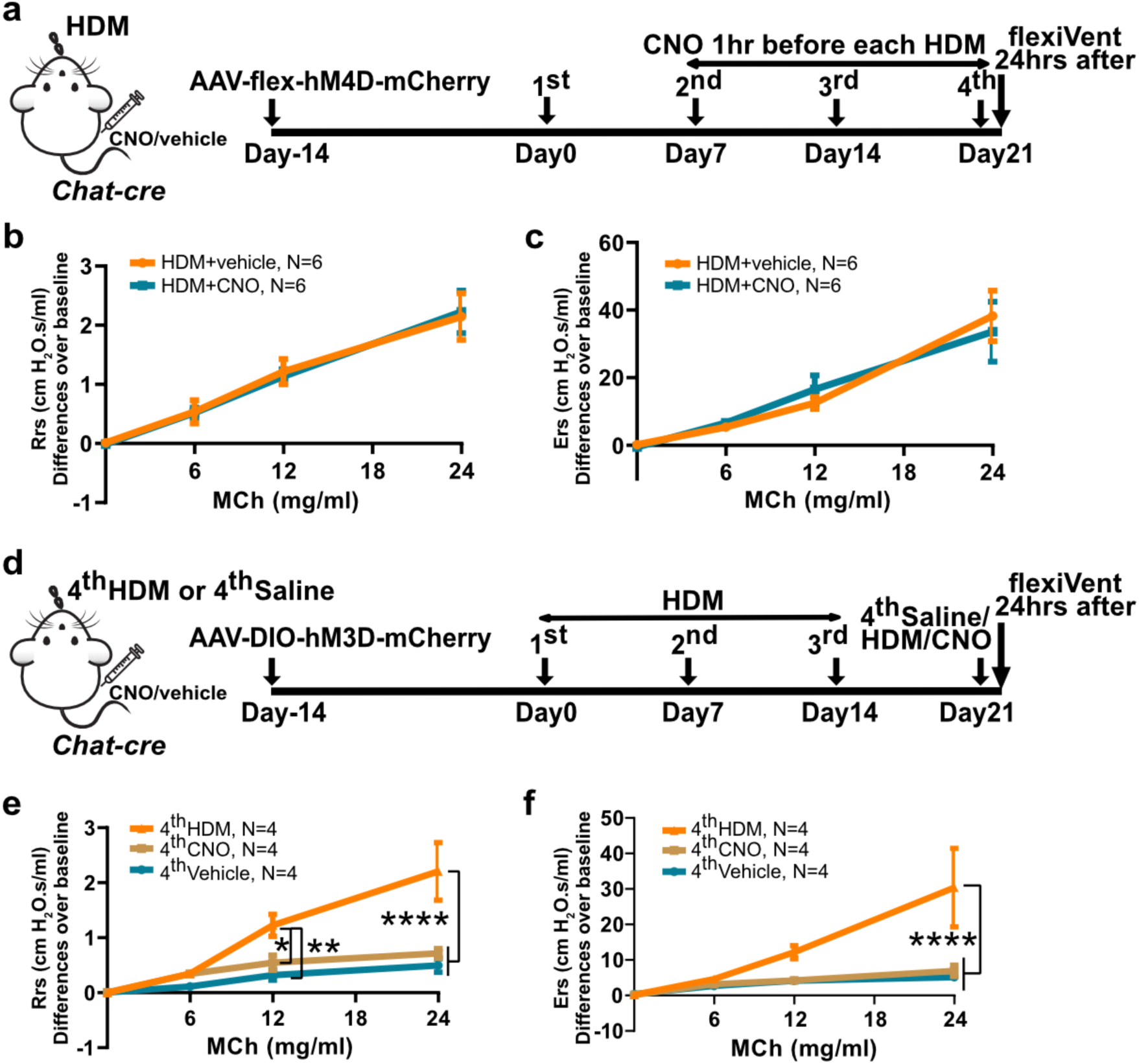
Parasympathetic neurons in the DMV or 12N did not mediate allergen-induced airway hyperreactivity. (**a**) Experiment scheme for chemogenetic inhibition of *Chat^+^* neurons in the DMV. (**b**, **c**) FlexiVent data showing no change in airway hyperreactivity after CNO injection. (**d**) Experiment scheme for chemogenetic activation of *Chat+* neurons in the DMV. (**e**, **f**) FlexiVent data showing no change in airway hyperreactivity after CNO injection, in place of the 4^th^ HDM. Two-way ANOVA was used for b, c, e and f, analysis was performed separately at each MCh concentration. Error bars represent means ± SEM. N.S., not significant, P ≥ 0.05.

**Extended Data Fig. 23.**
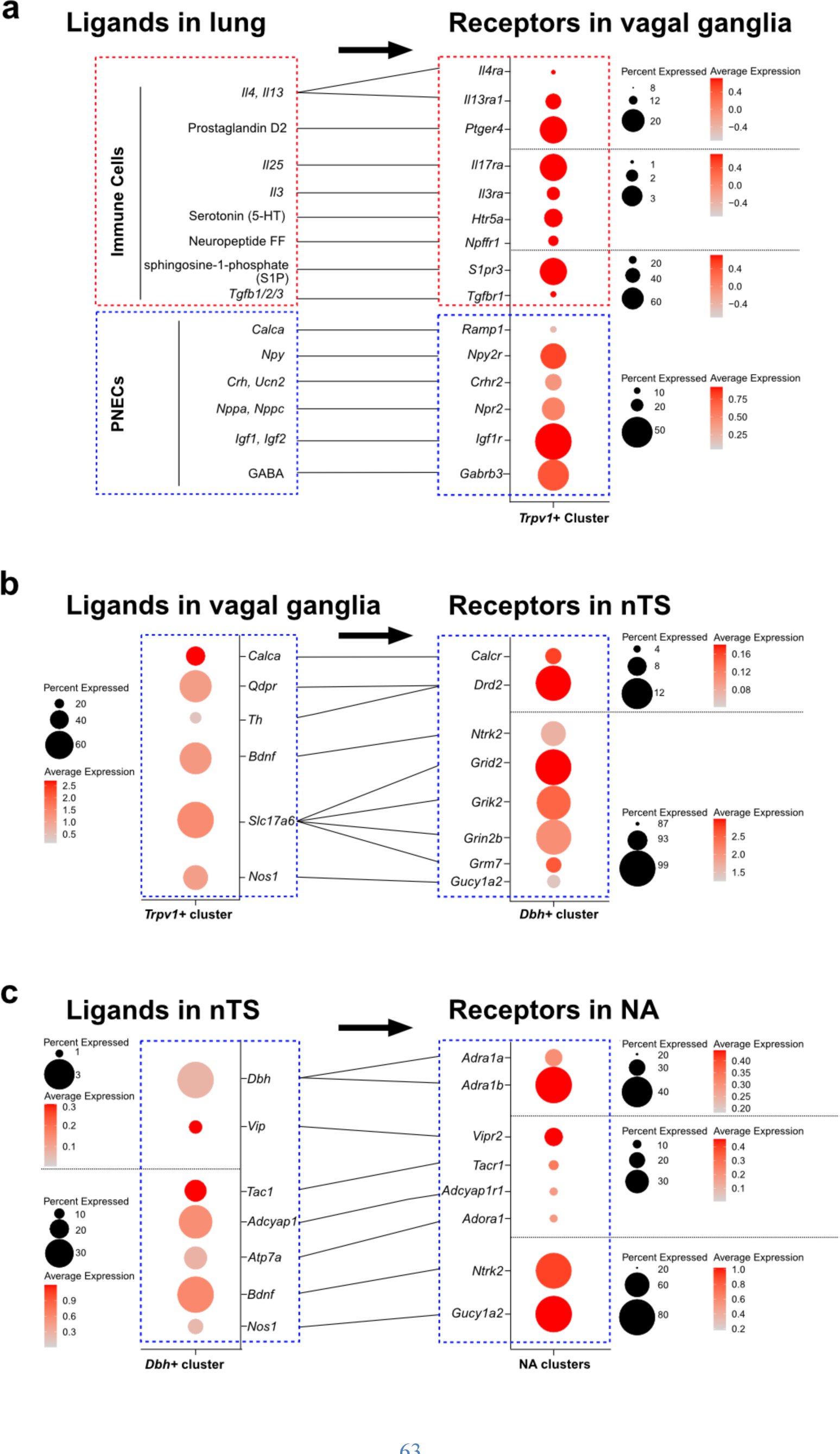
A predicted neuro-immune interaction network between the lung and the brainstem. (**a-c**) A Predicted neuron-immune interaction network between the lung, *Trpv1^+^*neurons in the vagal ganglia (a), *Dbh^+^* neurons in the nTS (b) and *Chat^+^*neurons in the NA (c) of the brainstem (connected by lines, arrows at top show the signaling direction). Red dashed lines outline genes participate in immune responses and blue dashed lines outline genes participate in neural signaling pathways.

**Extended Data Fig. 24.**
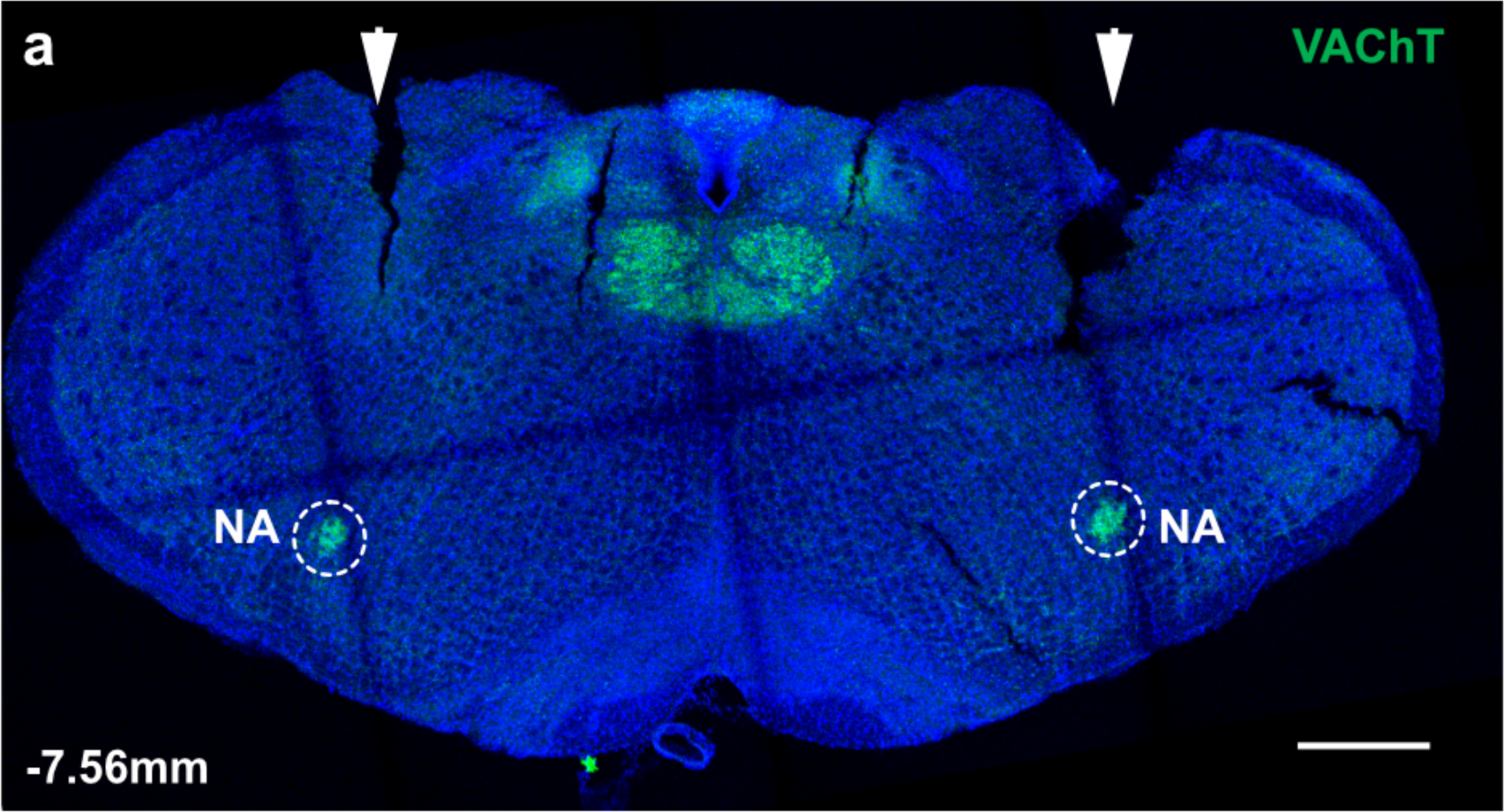
Cannula plantation in bilateral NA. (**a**) Representative cryosection (99 µm) of the NA showing bilateral cannula implantation to target NA. Arrowheads indicate the tract of cannula implantation. *Chat^+^* neurons in the NA (circled) were labeled with VAChT antibody. Scale bar, 0.5 mm.

